# Tracing the origin of pathologic pulmonary fibroblasts

**DOI:** 10.1101/2022.11.18.517147

**Authors:** Tatsuya Tsukui, Dean Sheppard

**Affiliations:** Division of Pulmonary, Critical Care, Allergy and Sleep, Department of Medicine, University of California, San Francisco, San Francisco, CA, USA

## Abstract

Fibroblasts substantially remodel extracellular matrix (ECM) in response to tissue injury and generate fibrotic scars in chronic diseases. Recent studies have identified diverse fibroblast subsets in healthy and injured tissues. However, the origin(s) and functional importance of injury-induced fibroblast lineages remain unclear. Here we show that alveolar fibroblasts, which provide a niche for maintaining alveolar type 2 cells in uninjured lungs, are the dominant source of multiple emergent fibroblast subsets that sequentially arise to facilitate fibrosis after lung injury. We demonstrate that Cthrc1+ fibroblasts, which express the highest levels of ECM proteins at injured sites, are effector cells for fibrogenesis using a novel mouse tool, Cthrc1-CreER. We use another novel mouse tool, Scube2-CreER, that uniquely targets alveolar fibroblasts, to reveal that alveolar fibroblasts are the dominant origin for multiple fibroblast subsets that emerge after lung injury. Pseudotime and in vitro analysis suggest that inflammatory cytokines initially induce chemokine-producing inflammatory fibroblasts from alveolar fibroblasts, which can differentiate into Cthrc1+ fibrotic fibroblasts in response to TGF-β. We identify similar fibroblast lineages from scRNA-seq in human pulmonary fibrosis. These results elucidate the pathologic fibroblast lineage development in response to lung injury and suggest that targeting key steps in transitions among these subsets could provide novel strategies for the treatment of fibrotic diseases.

## Main

Fibroblasts provide structural support to every organ by maintaining ECM architecture^1^. In response to tissue injury, new subsets of fibroblasts, including cells that produce large amounts of ECM, emerge at injured sites. These cells play an important role in normal tissue repair, but have been suggested to contribute to pathologic fibrosis in the setting of chronic diseases, an important contributor to organ failure^2,3^. These pro-fibrotic fibroblasts have been historically described as myofibroblasts based on increased expression of alpha-smooth muscle actin (α-SMA). We and others have recently shown that α-SMA is not specific to these pathologic ECM-producing fibroblasts and identified Cthrc1 (collagen triple helix repeat containing 1) as a more specific marker of the small subset of fibroblasts that produce the highest levels of ECM proteins in pulmonary fibrosis and other fibrotic diseases^4–7^. However, to date there has been no clear demonstration that Cthrc1+ fibroblasts actually contribute to fibrotic pathology. Furthermore, the origin of the cells that drive fibrotic pathology remains controversial. Although trans-differentiation from other cell types such as hematopoietic, epithelial, perivascular, and endothelial cells has been suggested^8,9^, there is little evidence from extensive scRNA-seq of fibrotic tissue to support these non-fibroblast sources. Recent scRNA-seq studies identified diverse fibroblast subsets in healthy tissues with distinct transcriptional profiles and anatomical locations^4,10^. Using computational lineage inference, one group recently proposed that pro-fibrotic fibroblasts likely universally emerge from adventitial fibroblasts, marked by high expression of the gene encoding peptidase inhibitor 16 (Pi16)^5^, while our previous computational lineage analysis suggested that pro-fibrotic fibroblasts in response to alveolar injury arise from alveolar fibroblasts^4^. Elucidating the origin(s) and trajectory for pro-fibrotic fibroblast development in response to tissue injury could lead to new insights into therapeutic targets for fibrotic diseases.

To experimentally determine whether Cthrc1+ fibroblasts contribute to fibrotic pathology and to determine the origin of these cells and how they emerge, we developed two novel tools that allowed functional assessment and lineage tracing. Using Cthrc1-CreER, we demonstrate the fibrogenic function of Cthrc1+ fibroblasts by ablating them in the bleomycin model of pulmonary fibrosis. Our previous work showed that signal peptide, CUB domain and EGF domain containing protein 2 (Scube2) is uniquely expressed in alveolar fibroblasts but not in other mesenchymal cells in mouse. Using Scube2-CreER mice, we confirm that alveolar fibroblasts provide a supportive niche for alveolar type 2 (AT2) cells in uninjured mice and demonstrate that they are the principal origin of multiple fibroblast subsets that emerge after lung injury, including pro-fibrotic Cthrc1+ cells. By lineage tracing with scRNA-seq we show that alveolar fibroblasts initially give rise to inflammatory fibroblasts after injury and that Cthrc1+ fibrotic fibroblasts emerge later. These fibroblast lineages are conserved in scRNA-seq data sets of human pulmonary fibrosis from multiple groups. Together, our data demonstrate that resident alveolar fibroblasts are the dominant source of all the fibroblast subsets that emerge in response to alveolar lung injury to drive post-injury fibrosis.

### Cthrc1+ fibroblasts are effector cells for fibrogenesis after lung injury

Our previous study showed Cthrc1+ fibroblasts uniquely emerge after lung injury, express the highest levels of ECM, and have enhanced migratory capacity^4^. In human idiopathic pulmonary fibrosis (IPF), CTHRC1+ fibroblasts are localized within fibroblastic foci, which are considered the leading edge of fibrogenesis^4,11^. To study the function of Cthrc1+ fibroblasts in lung injury, we generated Cthrc1-CreER mice by knocking a P2A-CreERT2-T2A-GFP sequence into the last exon of the Cthrc1 gene (Fig. 1a). Because GFP generated by this construct was not practically detectable by flow cytometry or tissue microscopy, we crossed Cthrc1-CreER mice with Rosa26-lox-stop-lox-tdTomato mice and injected tamoxifen on day 8 – 12 after inducing lung injury with intratracheal instillation of bleomycin (Fig. 1b). Whole lung imaging showed tdTomato+ cells emerged and formed aggregates in bleomycin-treated lungs (Fig. 1c). Flow cytometric analysis showed emergence of tdTomato+ cells among lineage-cells, which are negative for endothelial (CD31), hematopoietic (CD45), epithelial (EpCAM), and red blood cell (Ter119) markers, in bleomycin-treated lungs but not in saline-treated lungs (Fig. 1d, Supplementary Fig. 1a, b). tdTomato+ cells were closely associated with collagen 1 fibers on histology (Fig.1e). Quantitative PCR (qPCR) of purified tdTomato+ cells showed that expression of fibrotic genes such as collagen 1a1 (Col1a1), Cthrc1, and periostin (Postn) were highly enriched in tdTomato+ cells compared to all mesenchymal cells (lineage–cells) or all lung cells (Fig. 1f). Some tdTomato+ fibroblasts showed intermediate CD9 expression on day 14 (Supplementary Fig. 1c-e), consistent with our previous study^4^. Interestingly, tdTomato+ fibroblasts up-regulated CD9 on day 21 compared to day 14 (Supplementary Fig. 1c-e). These data suggest that Cthrc1-CreER successfully targets Cthrc1+ fibroblasts with the highest ECM expression after lung injury.

**Fig. 1.**
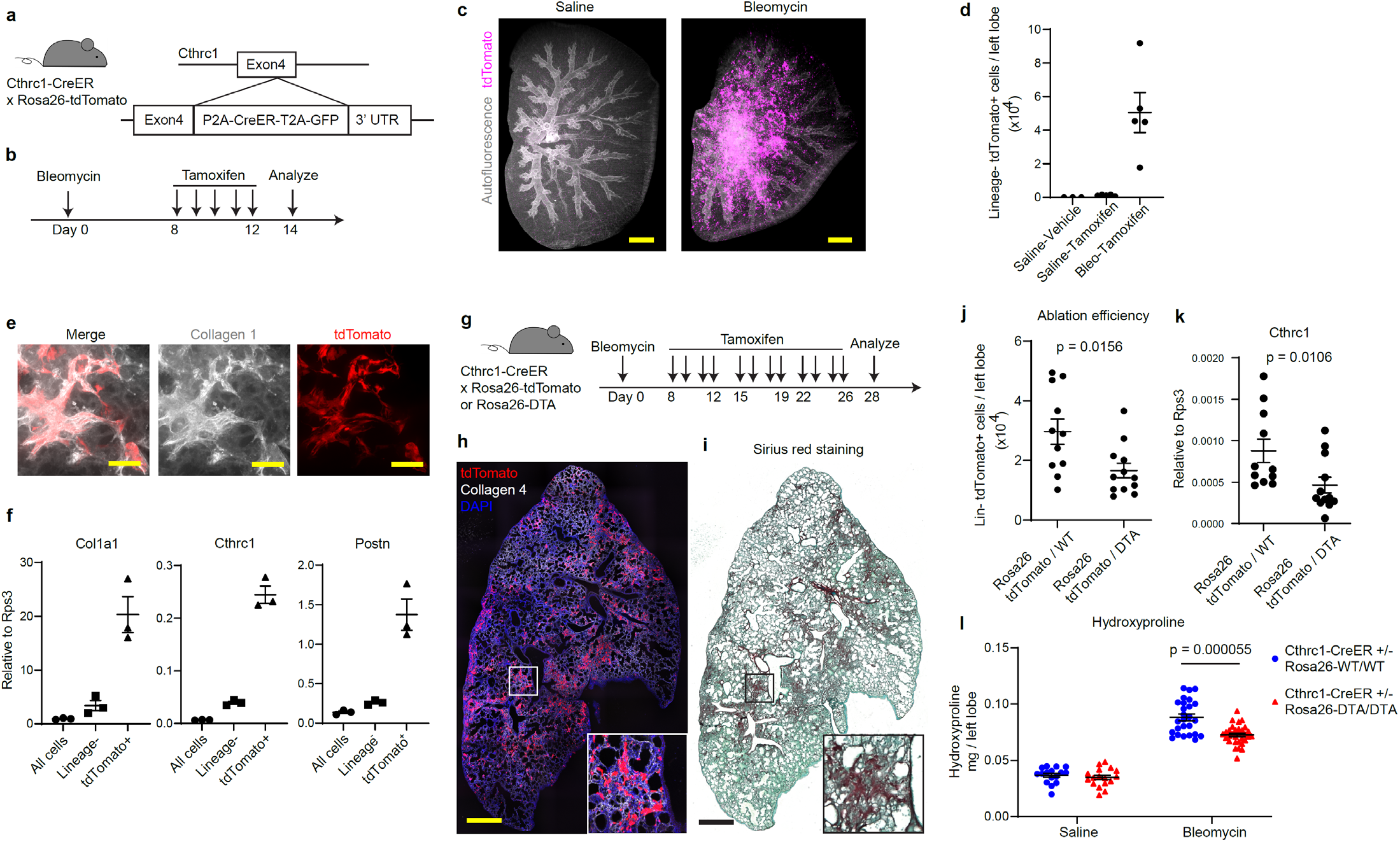
Cthrc1-CreER mouse demonstrates the pro-fibrotic function of Cthrc1+ fibroblasts. (a) Schematic of Cthrc1-CreER mouse generation. (b) Time course of bleomycin and tamoxifen treatment for day 14 analysis (c) Maximum projection of whole lung imaging. Autofluorescence is shown in grey. tdTomato is shown in magenta. (d) Flow cytometric cell count of lineage (CD31, CD45, EpCAM, Ter119)–tdTomato+ cells on day 14. n = 3 or 5 mice. (e) Collagen 1 staining (shown in grey) on day 14. tdTomato is shown in red. (f) qPCR analysis of all lung cells, lineage-cells, and tdTomato+ cells. Y axis is relative expression level compared to a house keeping gene Rps3. n = 3 mice. (g) Time course of bleomycin and tamoxifen treatment for day 28 analysis. (h) Fluorescence image of a lung section on day 28. Area of white square is magnified at the bottom right. tdTomato is shown in red. Collagen 4 staining is shown in grey. DAPI staining is shown in blue. (i) Sirius Red staining of a serial section of (h). Area of black square is magnified at the bottom right. (j, k) Ablation efficiency assessed by lineage–tdTomato+ cell number in left lobes (j) or by qPCR of whole lung cells for Cthrc1 (k) from Cthrc1-CreER+/- Rosa26-tdTomato/WT (n = 11 mice) or Cthrc1-CreER+/- Rosa26-tdTomato/DTA (n = 12 mice). (l) Hydroxyproline assay on day 28 of Cthrc1-CreER+/- Rosa26-WT/WT (n = 16 mice for saline, n = 26 mice for bleomycin) or Cthrc1-CreER+/- Rosa26-DTA/DTA (n = 17 mice for saline, n = 31 mice for bleomycin). Scale bars, 1 mm (c, h, i), 50 μm (e). Data are representative of at least two independent experiments except (l), which is a pool from two independent experiments. Data are mean ± SEM. Statistical analysis was performed using Mann-Whitney test (j, k, l).

We next used a tamoxifen injection protocol to label as many Cthrc1+ fibroblasts as possible until the peak of fibrosis (Fig. 1g). With this protocol, we observed close geographic association between tdTomato+ cells and fibrotic areas visualized by Sirius red staining on day 28 after bleomycin treatment (Fig. 1h, i). We then asked if ablation of Cthrc1+ cells by crossing with Rosa26-lox-stop-lox-DTA mice and using the same tamoxifen protocol reduces fibrosis. Approximately half of Cthrc1-CreER-labeled cells were killed by DTA induction (Fig. 1j, k). Despite this limited ablation efficiency, we observed a highly significant reduction in fibrosis measured by hydroxyproline content in the group with Cthrc1+ cell-ablation (Fig. 1l). Representative sections with Sirius red staining are shown in Supplementary Fig. 1f. These data demonstrate that Cthrc1+ fibroblasts are effector cells that significantly contribute to fibrosis after lung injury.

### Scube2-CreER-labeled alveolar fibroblasts are critical for maintenance of alveolar type 2 epithelial cells in uninjured lungs

We previously showed that healthy lungs contain multiple fibroblast subsets characterized by distinct anatomic localization in alveolar, adventitial or peribronchial regions^4^. Previously reported tools to label lung fibroblasts do not adequately distinguish among these populations. For example, Pdgfra and Tcf21 are broadly expressed in fibroblasts and inadequate to distinguish fibroblast subsets^4,5^. To specifically label alveolar fibroblasts, we generated Scube2-CreER mice (Fig. 2a). Scube2 is expressed in the clusters of fibroblasts we previously showed localize to the alveolar region but not in adventitial fibroblasts, peribronchial fibroblasts, pericytes, or smooth muscle cells^4^. We crossed Scube2-CreER mice with Rosa26-tdTomato mice and injected tamoxifen in the steady state (Fig. 2a). Flow cytometric analysis showed that 93.80 ± 0.73 % (n = 3, ± SEM) of tdTomato+ cells were negative for lineage markers (CD31, CD45, EpCAM, Ter119 and Mcam (smooth muscle cell and pericyte)), and that lineage-tdTomato+ cells were essentially all Seal–, CD9–, and Pdgfra+, which is consistent with our previous immunophenotyping of alveolar fibroblasts (Fig. 2b-g)^4^. After two weeks of tamoxifen treatment, labeling efficiency for lineage-, Sca1-, CD9-, and Pdgfra+ cells was 77.63 ± 3.34 % (n =3, ± SEM) (Supplementary Fig. 2a-b). We crossed Scube2-CreER/Rosa26-tdTomato mice with Col1a1-GFP (Col-GFP) reporter mice, where all fibroblasts express GFP, and checked tdTomato+ cell localization compared to GFP+ cells. By whole lung imaging, tdTomato+ signals were found diffusely throughout the alveolar region, while Col-GFP signals prominently highlighted bronchovascular bundles due to the higher cellular density of fibroblasts in adventitial and peribronchial locations (Fig. 2h). Imaging thick sections also confirmed the alveolar localization of tdTomato+ cells (Fig. 2i). Col-GFP+ cells in adventitial cuff spaces and peribronchial areas co-localized with the area of Pi16 staining and were not labeled by tdTomato (Supplementary Fig. 2c). These data validate that Scube2-CreER labels alveolar fibroblasts but not other fibroblast subsets.

**Fig. 2.**
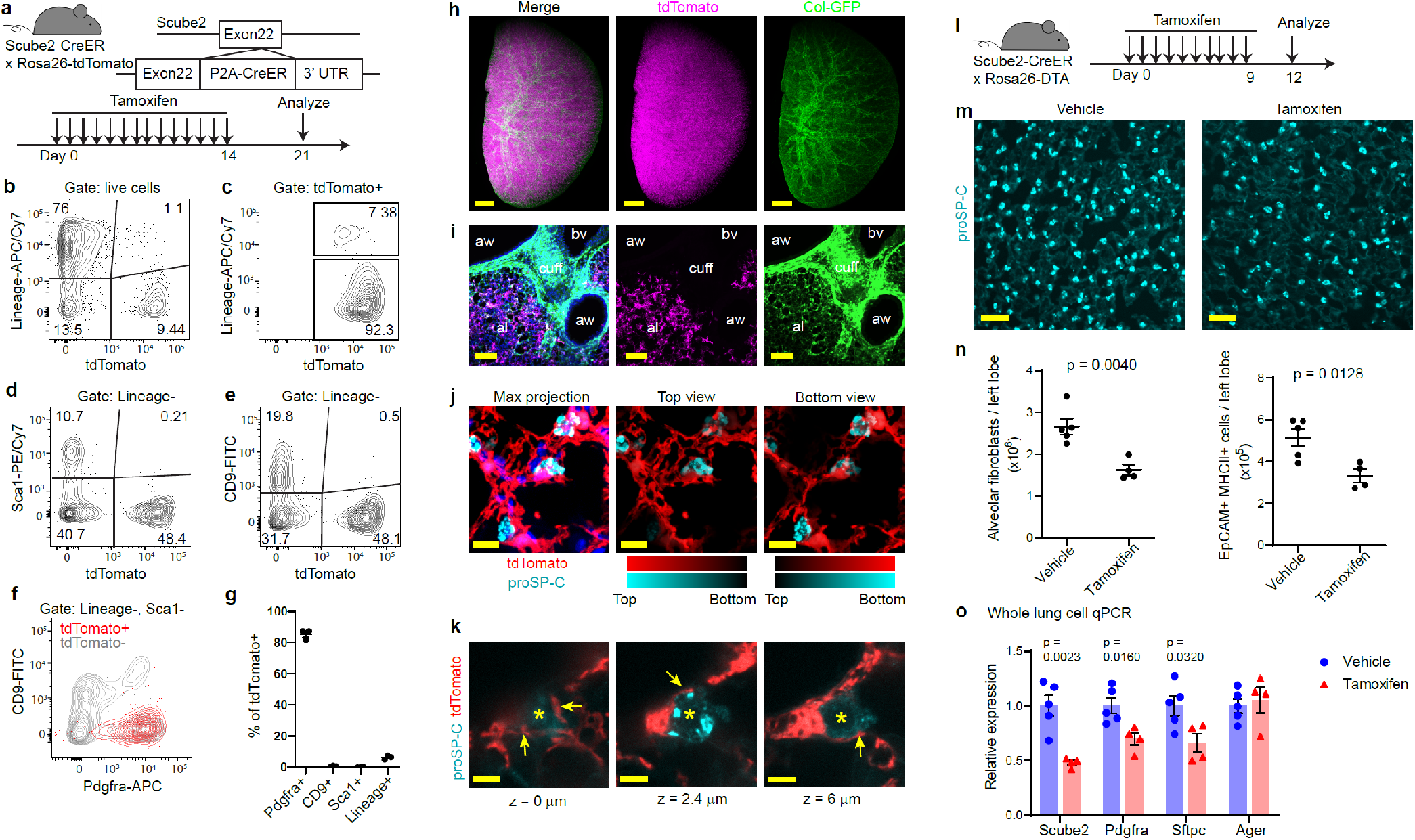
Scube2-CreER specifically labels alveolar fibroblasts and ablation of these cells decreases the numbers of alveolar type 2 cells in uninjured lungs. (a) Schematic of Scube2-CreER mouse generation and time course of tamoxifen treatment. (b-f) Flow cytometric analysis of lung cells from Scube2-CreER mouse. (g) Percent tdTomato+ in each marker+ fraction quantified by flow cytometry. Lineage markers include CD31, CD45, EpCAM, Ter119, and Mcam. Pdgfra+, CD9+, and Sca1+ fractions were pre-gated on lineage-cells. n = 3 mice. (h) Maximum projection of whole lung imaging. (i) Maximum projection of 32 z-stack images with step size 0.9 μm taken from a cleared thick section. aw, airway. bv, blood vessel. al, alveoli. cuff, cuff space. (j) 28 z-stack images with step size 0.5 μm shown as maximum projection (left), color-coded projection to the depth from top (middle), or color-coded projection to the depth from bottom (right). (k)Three representative planes from z-stack images with z-positions shown below the images. Asterisk indicates the same proSP-C+ cell. Arrows point to projections extending from alveolar fibroblasts (l) Time course of tamoxifen treatment. (m) Representative lung sections with proSP-C staining. (n) Flow cytometric counting for alveolar fibroblasts or AT2 cells gated as EpCAM+ MHCII+. n = 4 or 5 mice. (o) qPCR analysis of all lung cells. Y axis is relative expression level normalized to vehicle values. tdTomato is shown in magenta. Col-GFP is shown in green (h, i) DAPI staining is shown in blue (i, j). tdTomato is shown in red (j, k). ProSP-C staining is shown in cyan (j, k, m). Scale bars, 1 mm (h), 100 μm (i), 10 μm (j), 5 μm (k), 50 μm (m). Data are mean ± SEM. Data are representative of at least two independent experiments. Statistical analysis was performed using unpaired two-tailed t-test.

Previous work showed that Pdgfra+ lipofibroblasts in alveolar walls are closely associated with alveolar type 2 epithelial (AT2) cells, and Pdgfra+ cells isolated from lungs can support the growth of AT2 cells ex vivo^12,13^, suggesting that these cells might contribute to a supportive niche for AT2 cell maintenance. Our previous scRNA-seq data showed that the clusters of alveolar fibroblasts that express Scube2 also express a lipid droplet marker, Plin2^4^. We asked if Scube2-CreER-labeled alveolar fibroblasts might directly contact AT2 cells and support their maintenance in uninjured lungs. Three-dimensional imaging revealed that pro surfactant protein C (proSP-C)+ AT2 cells were closely localized with the cell bodies of tdTomato+ alveolar fibroblasts (Fig. 2j, k, Supplementary video 1). Those alveolar fibroblasts extended projections around AT2 cells (Fig. 2k). We crossed Scube2-CreER mice with Rosa26-DTA mice to see the impact of alveolar fibroblast ablation on AT2 cells (Fig. 2l). Three days after 10-day tamoxifen treatment, the frequency of proSP-C+ cells decreased on histology (Fig. 2m). Flow cytometric cell counting confirmed that the number of alveolar fibroblasts decreased 30 – 40% after ablation, and that AT2 cells identified as EpCAM+ MHC class II (MHCII)+ also decreased by 30-40% (Fig. 2n, Supplementary Fig. 2d-f)^14^. The decrease of alveolar fibroblast and AT2 cell markers was also confirmed by qPCR of whole lung (Fig. 2o), although no structural abnormality was observed at this early time point in the absence of injury (Supplementary Fig. 2g, h). These data provide in vivo evidence of a role for alveolar fibroblasts in maintaining a supportive niche for AT2 cells in the steady state.

### Lineage tracing by scRNA-seq identifies alveolar fibroblasts as the principal source of multiple emergent fibroblast subsets after lung injury

To investigate the fate of alveolar fibroblasts after lung injury, we performed scRNA-seq in Scube2-CreER/Rosa26-tdTomato mice. Alveolar fibroblasts were genetically labeled with tdTomato in the steady state by tamoxifen and lung injury was induced 2 weeks after the last tamoxifen injection (Fig.3a). We collected lungs on day 0 (untreated), 7, 14, and 21 with 3 biological replicates at each time point (Fig.3a). All mesenchymal populations were purified for scRNA-seq by magnetic and flow cytometric sorting for CD31-, CD45-, EpCAM-, and Ter119-cells, since our previous study showed that cells with high ECM expression are not marked by these lineage markers (Fig. 3a, Supplementary Fig. 3a)^4^. After processing and combining all the samples, we identified 11 clusters from 47,476 cells (Fig. 3b-e, Supplementary Fig. 3b-f). In addition to the mesenchymal subsets we found in our previous study (alveolar, adventitial, peribronchial, smooth muscle, mesothelial, and pericyte), we found 4 distinct clusters that emerged over this time course in response to lung injury. We labeled these as fibrotic, inflammatory, stress-activated, and proliferating based on the patterns of gene expression in each cluster (Supplementary Fig. 3c-f). Fibrotic fibroblasts were characterized by expression of Cthrc1 and high expression of Col1a1, Col3a1 and other components of the pathologic ECM, including tenascin C (Tnc) and Postn (Fig. 3d, e). Inflammatory fibroblasts expressed chemokines such as Cxcl12 and were marked by specific expression of serum amyloid A3 (Saa3) and lipocalin 2 (Lcn2) and interferon responsive genes, such as interferon induced transmembrane proteins, Ifitm2 and Ifitm3 (Fig. 3d, e). Gene ontology (GO) enrichment analysis of differentially expressed genes in inflammatory fibroblasts suggested responses to inflammatory cytokines including interferons, tumor necrosis factor (TNF) and interleukin 1 (IL-1) (Supplementary Fig. 3d, f), similar to the inflammatory fibroblast subsets reported in arthritis or cancer^15,16^. Stress-activated fibroblasts were characterized by the expression of the cell cycle arrest marker p21 (Cdkn1a), translation-related genes, and stress-related genes (dna damage-inducible transcript 3 (Ddit3) and bcl2-like protein 4 (Bax)) (Fig. 3d, e, Supplementary Fig. 3e, f). There were very few cells in any of these 4 subsets in the absence of injury on day 0 (Fig. 3c, Supplementary Fig. 3b). The frequency of inflammatory and proliferating fibroblasts peaked at day 7 and decreased at later time points (Fig. 3c). Fibrotic fibroblasts started to emerge on day 7 but their frequency increased on day 14 and 21 (Fig. 3c).

**Fig. 3.**
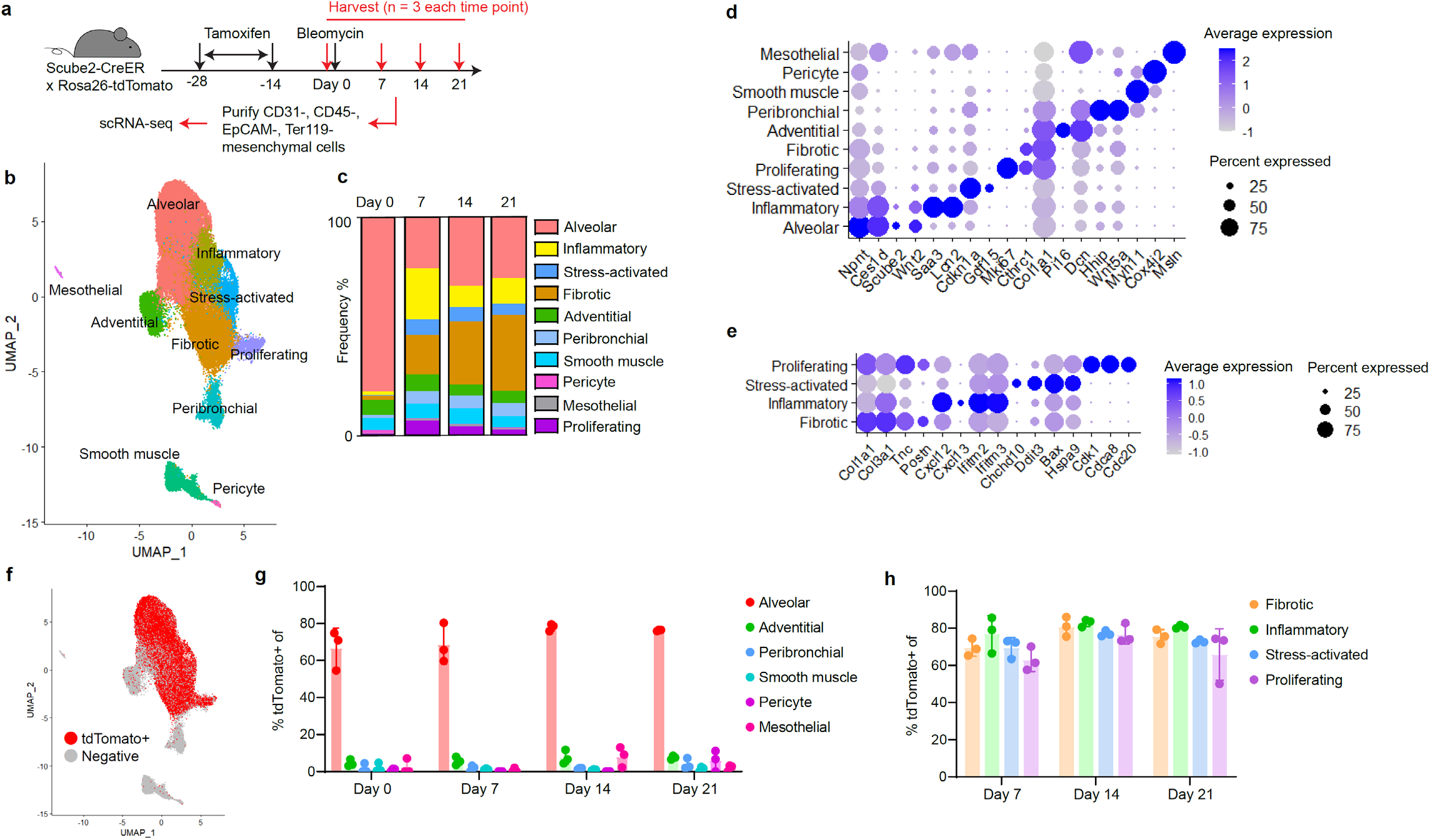
Lineage tracing by scRNA-seq reveals alveolar fibroblasts as origin of multiple emergent fibroblast subsets. (a) Schematic of scRNA-seq experimental design (b) UMAP plot of all scRNA-seq data. Clusters were shown with different colors. (c) Frequency of each cluster on different time points. (d) Dot plot showing representative markers for each cluster (e) Dot plot showing markers for subsets that emerge after injury. (f) UMAP plot showing tdTomato+ and tdTomato-negative cells. (g) Percent tdTomato+ of each subset that were present in normal lungs. (h) Percent tdTomato+ of each subset that emerged after injury. Data are mean ± SEM.

To assess the contribution of alveolar fibroblasts to each of these subsets, we analyzed tdTomato expression in our scRNA-seq data set. Although almost all cells showed at least low level tdTomato expression due to the baseline leak at the Rosa26 locus in the tdTomato reporter line, cells that underwent CreER-mediated recombination showed much higher tdTomato expression (Supplementary Fig. 4a, b). We defined cells with log-normalized tdTomato level above 3.5 as tdTomato+ cells, and less than or equal to 3.5 as tdTomato-negative cells (Fig. 3f, Supplementary Fig. 4b). We then quantified the frequency of tdTomato+ cell in each cluster at each time point (Fig. 3g, h). Among the subsets that were present in the steady state, 70 – 80% of alveolar fibroblasts were tdTomato+ while the other subsets contained very low numbers of tdTomato+ cells as expected (Fig. 3g). The tdTomato+ frequency for each of the 4 subsets that emerged after injury was virtually identical to the tdTomato+ frequency of alveolar fibroblasts (Fig. 3h). This pattern was consistent across all the replicates (Supplementary Fig. 4c). Moreover, when we compared tdTomato+ frequencies for alveolar fibroblasts and Cthrc1+ fibrotic fibroblasts in individual replicates, all were close to the line of identity (Supplementary Fig. 4d). These data suggest that virtually all the newly emergent fibroblasts originate from Scube2-CreER-labeled alveolar fibroblasts, and contribution to these emergent fibroblast subsets from other cells, including non-mesenchymal cells, is insignificant.

We next evaluated emergence of fibrotic and inflammatory fibroblasts from alveolar fibroblasts after lung injury. Whole lung imaging showed accumulation of Scube2-CreER-labeled cells in aggregates in central regions of the lungs 14 days after bleomycin treatment (Supplementary Fig. 5a, b). Some tdTomato+ cells up-regulated CD9 on day 21 after bleomycin treatment, as seen in Cthrc1-CreER-labeled fibrotic fibroblasts (Supplementary Fig. 5c, d). Purified tdTomato+ CD9+ cells on day 21 expressed higher fibrotic fibroblast markers and lower alveolar fibroblast markers compared to all tdTomato+ cells or tdTomato+ cells from untreated mice (Supplementary Fig. 5e), suggesting that Scube2-CreER-labeled alveolar fibroblasts differentiated into Cthrc1+ CD9+ fibrotic fibroblasts after lung injury. We also confirmed that some Scube2-CreER-labeled cells became inflammatory fibroblasts by staining for Saa3 (Supplementary Fig. 5f). We performed Saa3 immunostaining in Cthrc1-CreER/Rosa26-tdTomato mice on day 14 after bleomycin treatment to evaluate the spatial relationship between inflammatory and fibrotic fibroblasts. At this time point, fibrotic and inflammatory fibroblasts appeared to aggregate in adjacent but not overlapping regions, potentially reflecting distinct cytokine microenvironments (Supplementary Fig. 5g).

### Pseudotime analysis indicates sequential differentiation from alveolar to inflammatory and fibrotic fibroblasts

Although the presence of multiple emergent fibroblast subsets has been reported in other organs or pathologies^15–18^, the lineage relationships among these subsets has not been characterized well. To address this question, we focused on alveolar fibroblasts and the 3 largest emergent populations, inflammatory, stress activated, and fibrotic fibroblasts and performed pseudotime analysis by Monocle 3^19^ (Supplementary Fig. 6a). This analysis suggested that stress-activated fibroblasts arose from inflammatory fibroblasts and were potentially in a terminal state (Supplementary Fig. 6a). Since fibrotic fibroblasts seemed to be the other terminal state and most trajectories went through inflammatory fibroblasts (Supplementary Fig. 6a), we further focused on lineage relationships among alveolar, inflammatory, and fibrotic fibroblasts (Fig. 4a, b). We visualized the changes in representative markers along the pseudotime (Fig. 4c, d, Supplementary Fig. 6b, c) and found that alveolar fibroblast markers such as Pdgfra, Tcf21, Inmt, Npnt, and Ces1d gradually decreased along the pseudotime towards fibrotic fibroblasts (Fig. 4c, d, Supplementary Fig. 6c). Inflammatory fibroblast markers such as Saa3, Lcn2, Sod2, Hp, Cxcl12, and Sfrp1 were up-regulated in the middle of the pseudotime projection, but were down-regulated later, along with up-regulation of markers of fibrotic fibroblasts such as Cthrc1 and osteopontin (Spp1) (Fig. 4c, d, Supplementary Fig. 6c). Overlay of some of these markers on UMAP plots indicated heterogeneity within fibrotic fibroblasts, with Spp1 and Cthrc1 expressed in distinct cells on different points of the pseudotime projection (Fig. 4b, Supplementary Fig. 6c). Together, these data suggest that inflammatory fibroblasts are induced early after injury from alveolar fibroblasts and serve as an intermediate for the eventual emergence of fibrotic fibroblasts.

**Fig. 4.**
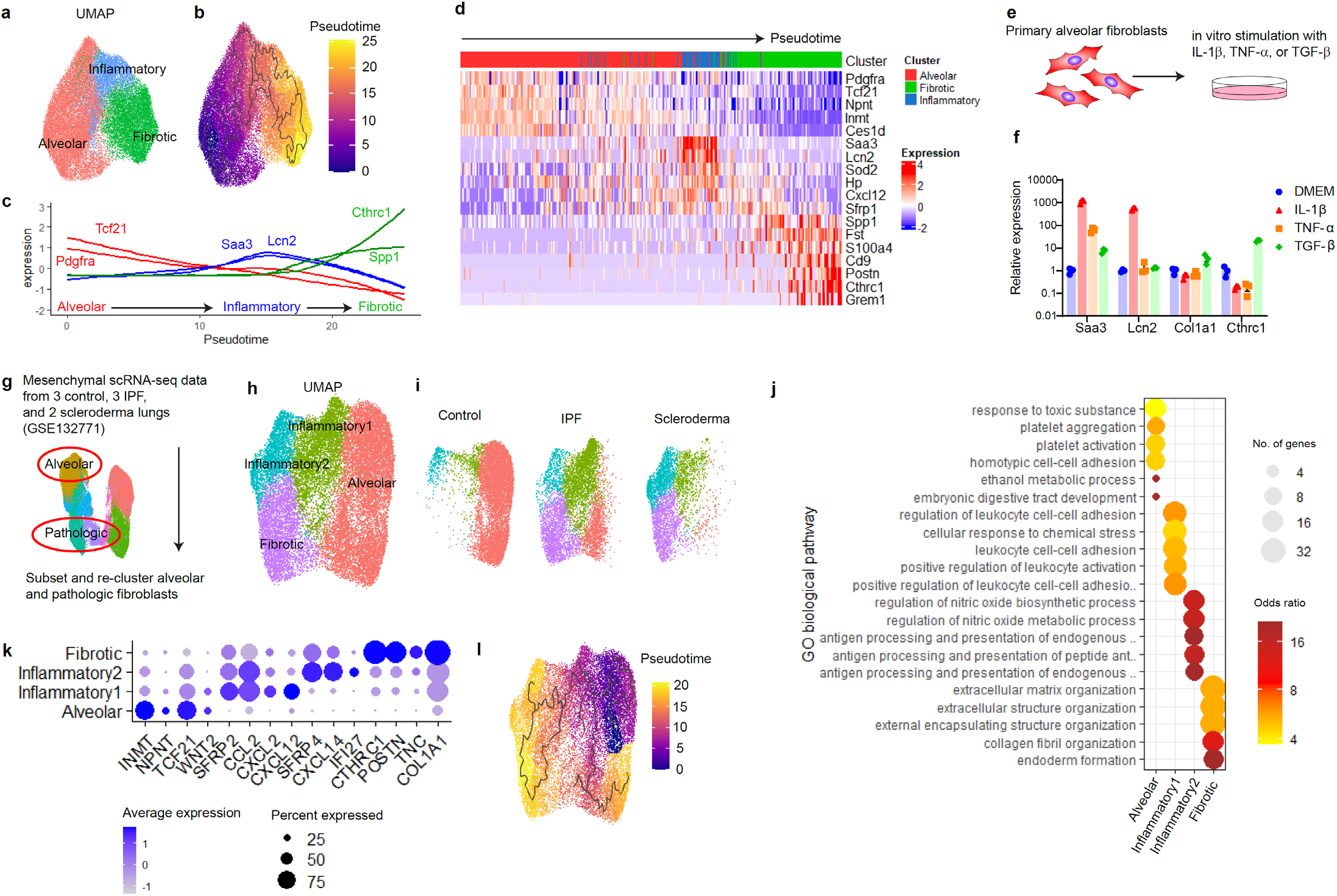
Alveolar fibroblasts sequentially differentiate into inflammatory and fibrotic fibroblasts in mouse and human pulmonary fibrosis. (a) UMAP plot of scRNA-seq data subsetted into alveolar, inflammatory, and fibrotic fibroblasts. (b) UMAP plot overlaid with pseudotime by Monocle 3. (c) Scaled expression of representative markers in pseudospace. (d) Heat map with cells arranged in pseudotemporal order showing changes in representative markers. Cluster annotations for each cell were shown above the heat map. (e, f) in vitro cytokine stimulation of primary alveolar fibroblasts. (e) Schematic of experiment (f) qPCR analysis for representative genes. Y axis is relative expression level normalized to DMEM (medium only) group. n = 3 wells. Data are mean ± SEM. Data are representative of three experiments. (g) Schematic of re-analysis of our previous human collagen-producing lung cell scRNA-seq data. (h) UMAP plot after subsetting and re-clustering alveolar and pathologic clusters. (i) UMAP plots for cells from control (n = 3), scleroderma (n = 2) or IPF (n = 3) lungs. (j) GO over-representation analysis for markers of each subset. (k) Dot plot for representative markers for each subset. (l) UMAP plot overlaid with pseudotime by Monocle 3. Arrow was manually drawn to show the direction of pseudotime.

### IL-1β and TGF-β1 are potent independent regulators of inflammatory and fibrotic fibroblasts

Since the differentially expressed genes of inflammatory fibroblasts indicated activation by inflammatory cytokines, we examined whether inflammatory cytokines could induce inflammatory fibroblast markers in alveolar fibroblasts in vitro (Fig. 4e). We cultured freshly isolated murine alveolar fibroblasts and stimulated them with IL-1β, TNF-α or TGF-β1 (Fig. 4e, f). We found that IL-1β dramatically up-regulated Saa3 and Lcn2 expression, while TGF-β1 up-regulated Col1a1 and Cthrc1 as previously described (Fig. 4f)^4^. We then asked if sequential treatment by IL-1β and TGF-β1 could induce fibrotic fibroblasts in vitro (Supplementary Fig. 6d). We found that TGF-β1 up-regulated Col1a1 and Cthrc1 within 24 hours regardless of prior stimulation by IL-1β (Supplementary Fig. 6e). In contrast, IL-1β-induced Saa3 and Lcn2 were down-regulated when cells were subsequently stimulated with TGF-β1 (Supplementary Fig. 6e). These data suggest that alveolar fibroblasts can rapidly differentiate into inflammatory or fibrotic fibroblasts in response to relevant exogenous stimuli and that TGF-β1 might serve as a critical switch during the transition of inflammatory fibroblasts to fibrotic fibroblasts.

### Alveolar-inflammatory-fibrotic fibroblast lineage relationship is also suggested in human pulmonary fibrosis

We next evaluated whether a similar relationship between alveolar, inflammatory, and fibrotic fibroblasts could be inferred from our own and others’ scRNA-seq data from human lungs. We first used our previously reported scRNA-seq data of collagen-producing lung cells from three patients with IPF, two with pulmonary fibrosis associated with scleroderma, and three control lungs^4^. We focused on alveolar and pathologic fibroblast clusters and re-clustered these to get higher resolution for heterogeneity (Fig. 4g-I, Supplementary Fig. 7a-d). We identified two previously unrecognized clusters expressing inflammatory chemokines in addition to the CTHRC1+ fibrotic cluster, which were enriched in IPF and scleroderma samples (Fig. 4h, i, k, Supplementary Fig. 7a, b). GO enrichment analysis suggested that inflammatory cluster 1 was potentially induced by IL-1 and/or TNF, while inflammatory cluster 2 was potentially induced by interferons (Supplementary Fig. 7c, d). Transcriptomic comparison to mouse emergent clusters also supported the similarity between mouse inflammatory cluster and human inflammatory cluster 1, as well as the similarity of fibrotic clusters and alveolar clusters from both species (Supplementary Fig. 7e). Although we did not observe a distinct cluster of stress-activated fibroblasts in humans, each of these inflammatory clusters were also enriched with GO terms related to stress responses such as “response to unfolded protein” (Supplementary Fig. 7c, d). Analysis of unique GO terms with Fisher test among all clusters showed features for antigen presentation in inflammatory cluster 2 (Fig. 4j), suggesting potential immune-modulating roles as has been recently described for a subset of cancer associated fibroblasts^20^. Pseudotime analysis by Monocle 3 showed a trajectory from alveolar fibroblasts through each inflammatory cluster toward the fibrotic cluster (Fig. 4i), consistent with the results from our mouse model.

We next examined if this lineage was observed in other data sets with larger sample size for human pulmonary fibrosis. We extracted alveolar and pathologic fibroblast clusters from Adams et al. and Habermann et al.^21,22^, and merged these cells with our alveolar and pathologic fibroblast clusters (Supplementary Fig. 8a-h). Based on the cluster annotation data transferred from Fig. 4h, we identified the two inflammatory, two alveolar, and fibrotic clusters in the merged data set (Supplementary Fig. 8e-f). Although the fibroblasts from our data outnumbered the fibroblasts from each of the other datasets probably due to the differences in the dissociation and processing protocols, we confirmed the presence of two inflammatory and fibrotic fibroblasts in multiple lung samples from Adams et al. and Habermann et al. (Supplementary Fig. 8f, g). These results suggest that alveolar-inflammatory-fibrotic fibroblast lineage we found in mouse lung injury is conserved in human pulmonary fibrosis.

## Discussion

Our data demonstrate that Cthrc1+ fibroblasts, which have been described to emerge in the settings of IPF^4^, scleroderma-associated pulmonary fibrosis^4,23^, SARS-CoV-2-associated lethal pneumonia^7^, myocardial infarction^6^, and cancer^5,24^, are important effector cells in driving tissue fibrosis. We also show that Scube2+ alveolar fibroblasts, which provide a niche for AT2 cell maintenance in the steady state, are the dominant source of all the fibroblast subsets including Cthrc1+ fibrotic fibroblasts that emerge in response to the alveolar fibrogenic injury we studied. Our GO enrichment analysis, pseudotime analysis, and in vitro studies suggest that alveolar fibroblasts are likely induced to differentiate into inflammatory fibroblasts by the effects of inflammatory cytokines during the initial phase of injury, and that fibrotic fibroblasts are later induced from inflammatory or alveolar fibroblasts by pro-fibrotic cytokines like TGF-β1. Importantly, analysis of scRNA-seq data from human pulmonary fibrosis suggests that similar fibroblast lineages are conserved in human pulmonary fibrosis. Although there are conflicting reports about the molecular function of Cthrc1 in acute injury and fibrosis^25,26^, our ablation experiments using Cthrc1-CreER demonstrate that Cthrc1+ fibroblasts themselves are important contributors to pulmonary fibrosis. Interestingly, the proportion of Cthrc1+ fibroblasts on day 21 in our scRNA-seq is only approximately 10% of all mesenchymal cells. Although the limitation in recombination and DTA-mediated killing efficiency resulted in only partial reduction in fibrosis after Cthrc1+ cell ablation, the significant effect of removing such a small population highlights the importance of Cthrc1+ fibroblasts in de novo fibrogenesis at injured sites. Since Cthrc1 is up-regulated in fibroblasts in multiple organs with fibrotic pathology, the Cthrc1-CreER line we describe here will be a useful tool to study fibrotic fibroblasts in many contexts.

Previous efforts to trace the pro-fibrotic fibroblast lineage in pulmonary fibrosis were limited by lack of understanding of the fibroblast subsets in the normal lung and those that emerge in the settings of lung injury and fibrosis. In this study, we developed a new mouse line, Scube2-CreER, that specifically distinguishes alveolar fibroblasts from adventitial fibroblasts, peribronchial fibroblasts, pericytes, and smooth muscle cells present in the normal lung. Using this line, we now clarify that adventitial fibroblasts, previously suggested based on computational analysis to be a universal source of profibrotic fibroblasts^5^, and other previously proposed progenitors such as pericytes, epithelial cells, endothelial cells, and hematopoietic cells are not important contributors to any of the new fibroblast subsets that emerge in the setting of fibrotic alveolar injury in the lung^8,9^, but rather that all of these subsets are principally derived from alveolar fibroblasts. Our results do not exclude the possibility, however, that other fibroblasts contribute to pathologic fibroblasts in other anatomic locations in the lung, such as perivascular or peri-airway fibrosis. Furthermore, since Scube2+ fibroblasts are uniquely present in the lung, it seems highly likely that Cthrc1+ pro-fibrotic fibroblasts that have been described in other fibrotic tissues emerge from different lineages. Further investigation of the mechanisms underlying the induction of each of these emergent populations and the functional contributions of each to progressive fibrosis or repair should lead to new therapeutic strategies targeting a wide array of diseases characterized by tissue inflammation and fibrosis.

## Supporting information

Supplementary Table 1 (qPCR primers)

## Abbreviations

ECM: extracellular matrix
IPF: idiopathic pulmonary fibrosis
Col-GFP: *Col1a1-EGFP*
α-SMA: α-smooth muscle actin
scRNA-seq: single-cell RNA-sequencing
AT2: alveolar type 2 epithelial
proSP-C: pro-surfactant protein C
GO: gene ontology
UMAP: Uniform Manifold Approximation and Projection

## Acknowledgments

We thank Dr. Junli Zhang at Gladstone Institute for support with generation of knockin mice and Dr. Walter Eckalbar at UCSF for support with computational analysis. T.T. was supported by Japan Society for the Promotion of Science (JSPS Overseas Research Fellowship), the Uehara Memorial Foundation, Mochida Memorial Foundation for Medical and Pharmaceutical Research, and the Frontiers in Medical Research Fellowship from the California Foundation for Molecular Biology. This work was supported by HL155786 (T.T.), HL142568 (D.S.), and a sponsored research agreement from AbbVie (D.S.). We thank UCSF core facilities: Laboratory for Cell Analysis supported by P30CA082103, Center for Advanced Light Microscopy supported by UCSF PBBR, Gladstone Transgenic Gene Targeting Core, Gladstone Genomics core, and Center for Advanced Technology supported by UCSF PBBR, RRP IMIA, and 1S10OD028511-01.

## Author contributions

T.T. and D.S. conceived the study, interpreted the data, and wrote the manuscript. T.T. performed and analyzed the experiments. D.S supervised the study.

## Methods

### Mice

Rosa26-lox-stop-lox-tdTomato (Stock No. 007914) and Rosa26-lox-stop-lox-DTA (Stock No. 009669) mice were obtained from the Jackson Laboratory. Col-GFP mice were obtained from Dr. David Brenner at University of California, San Diego^27^. Mice between the ages of 8 and 16 weeks old were used for the experiments. Male mice were used for Scube2-CreER scRNA-seq experiment. Both male and female mice were used in the other experiments. Heterozygous Cthrc1-CreER mice were used for experiments to avoid potential impact on fibrosis by altered Cthrc1 expression^25^. Homozygous Scube2-CreER mice were used for experiments to achieve higher recombination efficiency. No obvious phenotype of lung structure or fibrosis was observed in homozygous Scube2 CreER mice. For fibrosis induction, mice were treated with bleomycin in 75 μl saline by oropharyngeal aspiration. Since male mice develop more severe fibrosis, we used 2.5 U/kg bleomycin for male mice and 3 U/kg bleomycin for female mice, which were determined by induction of 7-9% body weight loss on day 7 and approximately 10% mortality rate. Male and female mice showed similar degree of fibrosis measured by hydroxyproline with these doses. Tamoxifen was dissolved in olive oil (Sigma) at 20 mg/ml, and 2 mg was intraperitoneally injected once a day. For labeling Cthrc1+ cells, tamoxifen was injected on day 8-12 after bleomycin treatment in most experiments. For ablating Cthrc1+ cells, tamoxifen was injected on day 8, 9, 11, 12, 15, 16, 18, 19, 22, 23, 25, and 26 after bleomycin treatment. Scube2-CreER mice were treated with tamoxifen for 2 weeks and used for experiment at least 1 week after the last tamoxifen injection unless specified. Heterozygous Cthrc1-CreER mice were used for experiments. Homozygous Scube2-CreER mice were used for experiments. Mice with homozygous Rosa26-tdTomato or Rosa26-DTA alleles were used for experiments unless specified. Mice were maintained in the UCSF specific pathogen–free animal facility in accordance with guidelines established by the Institutional Animal Care and Use Committee and Laboratory Animal Resource Center. All animal experiments were in accordance with protocols approved by the University of California, San Francisco Institutional Animal Care and Use Committee.

### Generation of Cthrc1-CreER and Scube2-CreER mice

The Cthrc1-CreER mouse strain was generated by homology-directed repair at the endogenous Cthrc1 locus aided by CRISPR/Cas9 endonuclease activity in C57BL/6 mice. Briefly, target sequence (5’-atatattggaatgccattac-3’), which had an adjacent PAM sequence, for guide RNA was selected to induce double strand breaks within the 3’UTR, and crRNA with input sequence GTAATGGCATTCCAATATAT and tracrRNA were obtained from IDT. A 2.38kb 5’homology arm was amplified from C57BL/6 mouse genomic DNA with forward primer 5’-GAGCTGAATGTTCAGGACCTCTTC-3’ and reverse primer 5’-TTTCGGTAGTTCTTCAATGATGAT-3’. A 2.15kb 3’ homology arm was amplified with forward primer 5’-CATTACAGTATTTAGTATTTCCTTCT-3’ and reverse primer 5’-ATTTGTTTGTTCCTAGGAGCTCTATAC-3’. A targeting vector with P2A-CreERT2-T2A-GFP-stop codon-rabbit beta globin polyA sequence flanked by 5’ and 3’ homology arms was generated using NEBuilder HiFi DNA Assembly (NEB) and cloned into a pKO2 backbone plasmid. The targeting vector was linearized at SalI (NEB) and NotI (NEB) sites flanking the donor DNA sequence and the linearized donor DNA was purified by agarose gel electrophoresis with GeneJet Gel Extraction kit (Thermo Fisher). Linearized donor DNA and CRISPR/Cas9 complex were injected into C57BL/6 fertilized zygotes, which were then implanted into oviducts of pseudopregnant female mice. 215 embryos were implanted and 17 pups were born. Three founders were identified by genotyping. We used one founder to expand the colony. The Scube2-CreER mouse strain was generated by a similar process. Target sequence of guide RNA for endogenous Scube2 3’UTR locus was 5’-GTGACTCGTCAGAGTTCAGT-3’ and input sequence for crRNA was ACTGAACTCTGACGAGTCAC. A 2.79 kb 5’homology arm was amplified with forward primer 5’-TGGCCTTGACTGTGTACACTTACATTA-3’ and reverse primer 5’-TTTGTAAGGCCTCAGAAACCTTGACACTTT-3’. A 2.24kb 3’homology arm was amplified with forward primer 5’-TTTTATAGACAATACAGATATCTTGA-3’ and reverse primer 5’-TGTGTGAGAATACATGTGTACCACA-3’. A targeting vector with P2A-CreERT2-stop codon-rabbit beta globin polyA sequence flanked by 5’ and 3’ homology arms was generated and linearized for injection. 220 embryos were implanted and 22 pups were born. 5 founders were identified and we used one of them to expand the colony.

### Tissue dissociation

Mouse lungs were harvested after perfusion through the right ventricle with 5 ml PBS. After mincing with scissors, the tissue was suspended in protease solution [0.25 % Collagenase A (Millipore Sigma), 1 U/ml Dispase II (Millipore Sigma), 2000 U/ml Dnase I (Millipore Sigma) in Hanks’ Balanced Salt Solution (Thermo Fisher)]. The suspension was incubated at 37°C for 60 min with trituration by micropipette every 20 min. Then the cells were passed through a 100 μm cell strainer (BD Biosciences), washed with PBS, and suspended in PBS with 0.5 % bovine serum albumin (BSA) (Fisher BioReagents).

### Flow cytometry

After tissue dissociation, 1 x 10^6^ cells were used for flow cytometry. Cells were resuspended in PBS with 0.5% BSA containing antibodies. For identifying lineage+ cells, cells were first stained with biotin-labeled antibodies for lineage markers, followed by washing and staining with other antibodies and streptavidin-A488 or APC/Cy7. DAPI (Thermo Fisher) was used at 0.1 μg/ml to identify dead cells. Flow cytometric cell count was performed using CountBright Plus Absolute Counting Beads (Invitrogen). The following antibodies were used at 1:200 unless specified: anti-CD9 (clone MZ3, FITC, APC/Fire750, biotin; BioLegend), anti-CD31 (clone 390, A488, biotin; BioLegend), anti-CD45 (clone 30F-11, PE/Cy7, biotin; BioLegend), anti-Mcam (clone ME-9F1, biotin; BioLegend), anti-Pdgfra (clone APA5, APC; BioLegend), anti-Epcam (clone G8.8, PE, biotin; BioLegend), anti-I-A/I-E (MHC class II) (clone M5/114.15.2, APC/Cy7; BioLegend) anti-Sca1 (clone D7, PE/Cy7, biotin; BioLegend), streptavidin-APC/Cy7 (BioLegend), streptavidin-A488 (1:1000, Thermo Fisher). Data acquisition or cell sorting was performed using FACS Aria III or Aria Fusion (BD Biosciences). Flow cytometry data were analyzed using FlowJo v10 (Becton Dickinson).

### Hydroxyproline assay

Fibrosis after bleomycin treatment was assessed by hydroxyproline assay of tissue lysates as described previously^28^. Briefly, left lobes were homogenized and precipitated with trichloroacetic acid. Following baking at 110°C overnight in HCl, samples were reconstituted in water, and hydroxyproline content was measured by a colorimetric chloramine T assay.

### scRNA-seq library preparation and sequencing

Scube2-CreER, Rosa26-tdTomato double homozygous mice were treated with tamoxifen for two weeks. Bleomycin treatment was performed 2 weeks after the last tamoxifen treatment. Three biological replicates from day 0 (non-bleomycin-treated), 7, 14, and 21 samples were collected on the same day and tamoxifen/bleomycin treatment was scheduled accordingly. After harvesting and dissociating left lobes, mesenchymal cells were enriched by magnetic negative selection with anti-CD31, CD45, EpCAM and Ter119-biotin antibodies (1:200) and Dynabeads MyOne Streptavidin T1 (40 μl / sample, Invitrogen). After magnetic negative selection, cells were stained with Streptavidin-APC/Cy7 (1:200) and DAPI (0.1 μg/ml). Approximately 2 x 10^5^ lineage-APC/Cy7-negative cells were sorted for each sample. The sorted cells were counted and labeled with oligonucleotide tags for multiplexing using 10x Genomics 3’ CellPlex Kit Set A. Tag assignment was as follows; day 0 (301, 302, 303), day 7 (304, 305, 306), day 14 (307, 308, 309), and day 21 (310, 311, 312). All 12 samples were pooled and 30,000 cells / lane were loaded onto 4 lanes of Chromium Next GEM Chip (10x Genomics). Chromium Single Cell 3’ v3.1 (10x Genomics) reagents were used for library preparation according to the manufacturer’s protocol. The libraries were sequenced on an Illumina NovaSeq 6000 S4 flow cell.

### Sequencing data processing

Fastq files were uploaded to the 10x Genomics Cloud Analysis website (https://www.10xgenomics.com/products/cloud-analysis) and reads were aligned to a custom reference of mouse genome mm10 with tdTomato-WPRE-polyA transcript sequence using Cell Ranger version 6.1.1. tdTomato-WPRE-polyA sequence was obtained from the sequence of the targeting vector for Ai9 mouse (Addgene plasmid #22799) since the Ai9 mouse shares the same sequence for tdTomato-WPRE-polyA with the Ai14 mouse used in this study^29^. The data were demultiplexed and multiplets identified by the presence of multiple oligonucleotide tags were removed using 10x Genomics cloud analysis function with default parameters. Raw count matrices were imported to the R package Seurat v4.1.0^30^ and cells with fewer than 200 detected genes, larger than 7500 detected genes, or larger than 15% percent mitochondria genes were excluded. We used the DoubletFinder package^31^ for individual samples to remove doublets that were not detected upon alignment using estimated multiple rate 2%. We then merged all the sample objects, identified the top variable genes using the Seurat implementation *FindVariableGenes*, and integrated the samples using the *RunFastMNN*^32^ function of the SeuratWrappers R package. For visualization, the *RunUMAP* function of Seurat was performed using MNN dimensional reduction. Nineteen clusters were initially identified using *FindNeighbors* and *FindClusters* functions of Seurat with resolution = 0.8 from a total of 47,809 cells. Cluster 17 (168 cells) was a cluster mixed with small number of lineage (CD31, CD45, and EpCAM)+ cells that were not removed by FACS sorting. Cluster 18 (165 cells) showed up in two different locations on the UMAP embedding, one close to alveolar fibroblasts and the other close to peribronchial fibroblasts. Cluster 18 cells expressed both alveolar and peribronchial fibroblast markers, suggesting that they were doublets that were not removed by prior processing. We excluded cluster 17 and 18, and re-clustered the remaining 47,476 cells with *FindVariableGenes, RunFastMNN, RunUMAP, FindNeighbors*, and *FindClusters* functions with clustering resolution = 0.3. Differentially expressed genes for each cluster were identified using the *FindAllMarkers* function of Seurat focusing on genes expressed by more than 25% of cells (either within or outside of a cluster) and with a log fold change greater than 0.25. tdTomato+ cells were defined by natural log-normalized tdTomato expression level greater than 3.5. The meta data including cluster, sample, and tdTomato+ annotations were exported for quantifying the tdTomato+ frequency in each cluster. Alveolar1 and Alveolar 2 clusters were combined as “Alveolar” in quantification. Gene ontology enrichment analysis for the differentially expressed genes was performed using DAVID (Database for Annotation, Visualization and Integrated Discovery) Bioinformatic Resources software version 2021, or using one-sided Fisher’s exact tests implemented in gsfisher R package (https://github.com/sansomlab/gsfisher/). We performed pseudotime analysis on the UMAP embeddings using Monocle 3 v1.0.0^19^, specifying cells on day 0 as roots of the pseudotime. Scaled expression of representative markers along the pseudotime was visualized using ggplot2 v3.3.6 (Fig. 4c). A heatmap with cells arranged in pseudotemporal ordering (Fig. 4d) was generated using Slingshot v2.2.0^33^ and ComplexHeatmap R package 2.10.0, specifying starting cluster as “Alveolar” and ending cluster as “Fibrotic”.

### Human scRNA-seq data processing

We used our previous human scRNA-seq data set of pulmonary fibrosis (GSE132771)^4^. We subsetted alveolar and pathologic fibroblast clusters from our mesenchymal cell data, and re-clustered them using *FindVariableGenes, RunFastMNN, RunUMAP, FindNeighbors*, and *FindClusters* functions of Seurat with clustering resolution = 0.3. Cluster markers were identified using *FindAllMarkers* function of Seurat with min.pct = 0.25 and logfc.threshold = 0.25. For comparison between human and mouse emergent clusters, average expression of the clusters was exported from scaled data of Seurat objects, and human genes were converted to mouse orthologs using biomaRt R package, followed by calculation of Spearman’s correlation coefficient by *cor* function of R. Pseudotime analysis was performed on the UMAP embeddings using Monocle 3, specifying cells from control lungs as roots of the pseudotime. For integrating alveolar and pathologic fibroblasts from Adams et al.^21^ and Habermann et al.^22^, we obtained their data sets from GSE147066 and GSE135893, respectively. Raw count matrix of mesenchymal cells of control and IPF lungs from Adams et al. was batch-corrected using *RunFastMNN* function of Seurat, and visualized by *RunUMAP, FindNeighbors*, and *FindClusters* functions of Seurat. Alveolar and pathologic fibroblast clusters were identified by examining markers such as INMT, NPNT, TCF21, CTHRC1, COL1A1, and POSTN. For Habermann et al., mesenchymal cells annotated by the original authors were subsetted from the Seurat object containing all cells. By examining markers for alveolar and pathologic fibroblasts, a cluster the authors annotated as myofibroblasts was identified as cells containing both alveolar and pathologic fibroblasts. We subsetted those alveolar and pathologic fibroblast clusters from Adams et al. and Habermann et al. data sets, merged all with our alveolar and pathologic fibroblast clusters, and integrated these data sets using *RunFastMNN* function of Seurat by splitting the object by individual patients or donors. After UMAP visualization and clustering, there were two minor clusters of which cells originated only from Adams et al.. One of these clusters was characterized by unusually high numbers of genes and read counts. The other cluster was characterized by high mitochondrial gene proportions. Since these two clusters were only seen in Adams et al. and seemed to be driven by technical artefacts but the other clusters from Adams et al. merged well with the other two data sets, we excluded these two clusters. We re-clustered the remaining cells and annotated the clusters based on the overlap with cells from our data set, which had transferred cluster annotations as shown in Fig. 4h.

### Histology, immunohistochemistry, and imaging

For histology, lungs were inflated with 4% PFA and immersed in 4% PFA over night at 4°C. The lungs were then immersed in 30% sucrose for 24 hours at room temperature, and then embedded in OCT. 12 μm sections for thin section histology or 100 μm sections for thick section histology were made using a cryostat CM1850 (Leica). Thin sections were attached to Superfrost Plus microscope slides (Fisher). For Sirius Red staining, sections were incubated with 0.1% Sirius Red in Saturated Picric Acid (Electron Microscopy Sciences) with 10% w/v Fast Green FCF (Fisher) for 1 hour. Thick sections were processed as floating sections in buffers. Thick sections were cleared using a CUBIC method^34^. After delipidation with CUBIC-L (TCI), sections were stained with anti-Pi16 (5 μg/ml, R&D, AF4929), anti-proSP-C (1:5000, Sigma-Aldrich, AB3786) anti-collagen 1 (1:200, Southern Biotech, 1310-01), anti-collagen 4 (1:5000, LSL, LSL-LB-1403), or anti-Saa3 (1:100, Abcam, JOR110A) followed by donkey anti-rabbit IgG-alexa 488 or 647 (1:1000, Thermo Fisher, A-21206, A-21245), donkey anti-goat IgG-alexa 647 (1:1000, Thermo Fisher, A-21447), or donkey anti-rat IgG-alexa 647 (1:1000, Thermo Fisher, A78947). Sections were then treated with CUBIC-R+(M) (TCI), placed in a well of glass bottom plate with sections covered with CUBIC-R+(M), and imaged by an inverted Crest LFOV spinning disk confocal microscope (Nikon Ti2). Images were processed using Image J version 1.53q. 3D-reconstruction of z-stack images was performed using Icy version 2.0. For whole lung imaging, 4% PFA-fixed lungs were cleared with CUBIC-L (TCI) and treated with CUBIC-R+(M), followed by imaging with Mounting Solution (RI 1.520, TCI) using a Nikon AZ100 microscope configured for light sheet microscopy. Autofluorescence signal in the GFP channel was used to visualize the lung structure except Fig. 2h. Maximum projection images were generated using Image J.

### Quantitative Real-Time PCR analysis

Approximately 2000 cells were directly sorted into TRIzol reagent (Thermo Fisher), and RNA was isolated according to manufacturer’s protocol. The RNA was reverse-transcribed using a Super Script IV VILO Master Mix with ezDNase Enzyme kit (Thermo Fisher). Quantitative Real-Time PCR was performed using PowerUp SYBR Green Master Mix (Thermo Fisher) with a Quant Studio 4 (Applied Biosystems). Primer sequences are listed in Supplementary Table 1.

### *In vitro* stimulation of primary alveolar fibroblasts

Alveolar fibroblasts were enriched by magnetic negative selection for CD31, CD45, EpCAM, Ter119, Sca1, and CD9. 2 x 10^5^ cells were seeded into 48 well plates and initially cultured in DMEM (Corning) with 2% fetal bovine serum (FBS) (Gibco) and 1% penicillin-streptomycin (Gibco) for 24 hr. Then medium was changed to serum-free DMEM with 1% penicillin/streptomycin for 24 hr. After serum starvation, medium was changed to serum-free DMEM with 1% penicillin/streptomycin, containing 1 ng/ml IL-1β, 1 ng/ml TGF-β1 (R&D, 7754-BH), or 10 ng/ml TNF-α (R&D, 210-TA). For sequential stimulation, the medium was changed to serum-free DMEM with 1% penicillin/streptomycin containing 1 ng/ml IL-1β, 1 ng/ml TGF-β1, or both for 24 hr. After the cytokine stimulations, cells were lysed by directly adding 400 μl Trizol into the wells. Cell culture was performed under standard conditions (37 °C, 5% CO_2_).

### Data analysis

Mean linear intercept was quantified as described previously^35^. scRNA-seq data analysis was performed in R version 4.1.3. Statistical tests were performed in GraphPad Prism version 9.4.0.

### Data availability

The scRNA-seq data generated in this study are deposited in Gene Expression Omnibus (GEO) (accession number will be released upon acceptance). The Seurat objects used in this study including re-analysis of publicly available human scRNA-seq data are available from the authors upon request.

### Code availability

The codes used in scRNA-seq analysis are available at GitHub (https://github.com/TatsuyaTsukui/AlveolarLineage).

---

**Supplementary Fig. 1.**
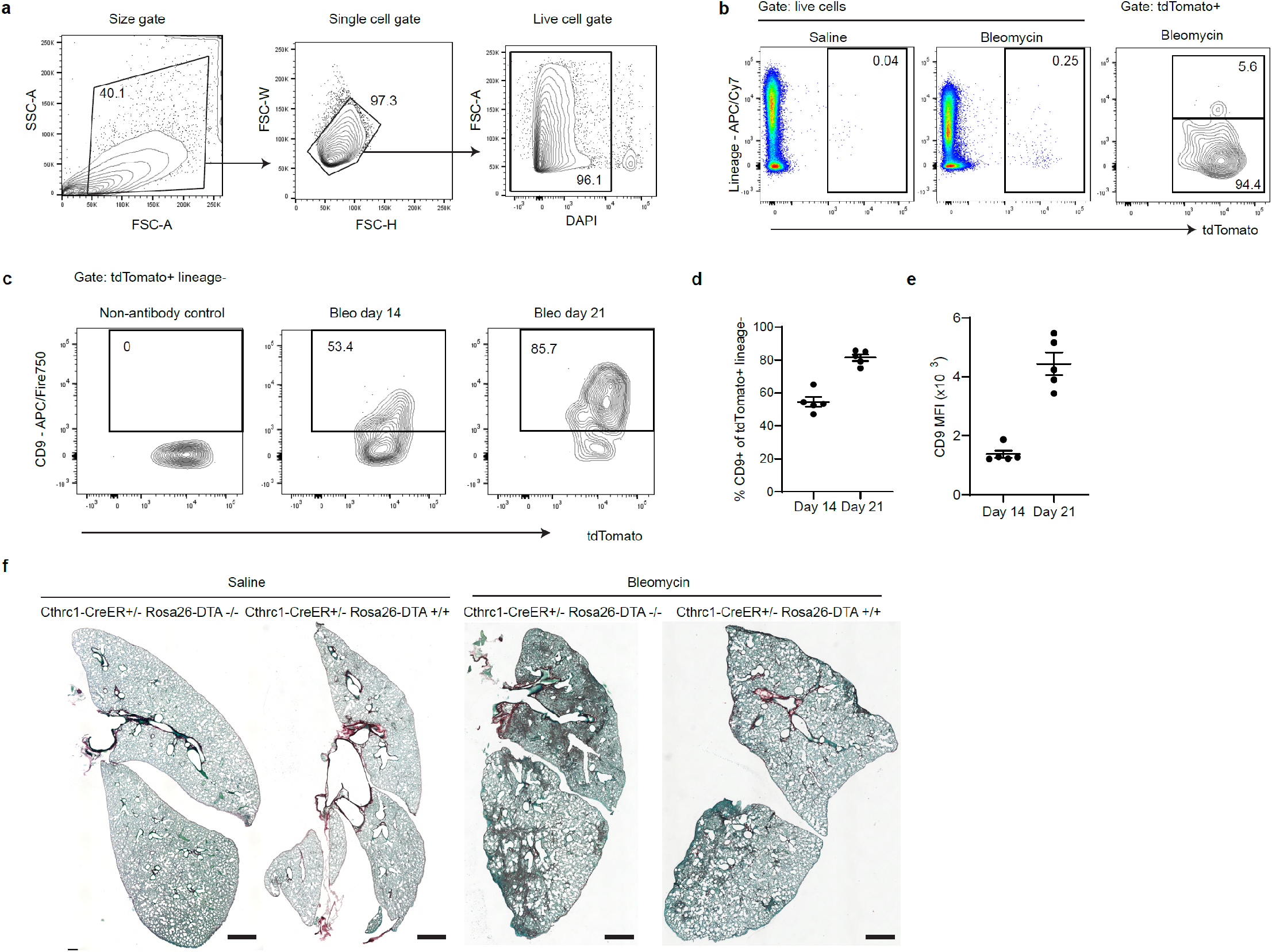
Cthrc1-CreER mouse demonstrates the pro-fibrotic function of Cthrc1+ fibroblasts. (a) Gating strategy for cell size, singlet, and live cells. (b) Flow cytometry plots show increase in lineage-tdTomato+ cells on day 14 after bleomycin treatment. (c) Flow cytometry plots show CD9 expression on tdTomato+ lineage-cells increases between day 14 and 21. (d) Flow cytometric quantification of percent CD9+ of tdTomato+ lineage-cells. n = 5 mice (e) Mean fluorescence intensity (MFI) of CD9 on tdTomato+ lineage-cells. n = 5 mice. (f) Representative images of Sirius Red staining of lung sections from the ablation experiment. Data are mean ± SEM. Data are representative of at least two independent experiments. Scale bars, 1 mm.

**Supplementary Fig. 2.**
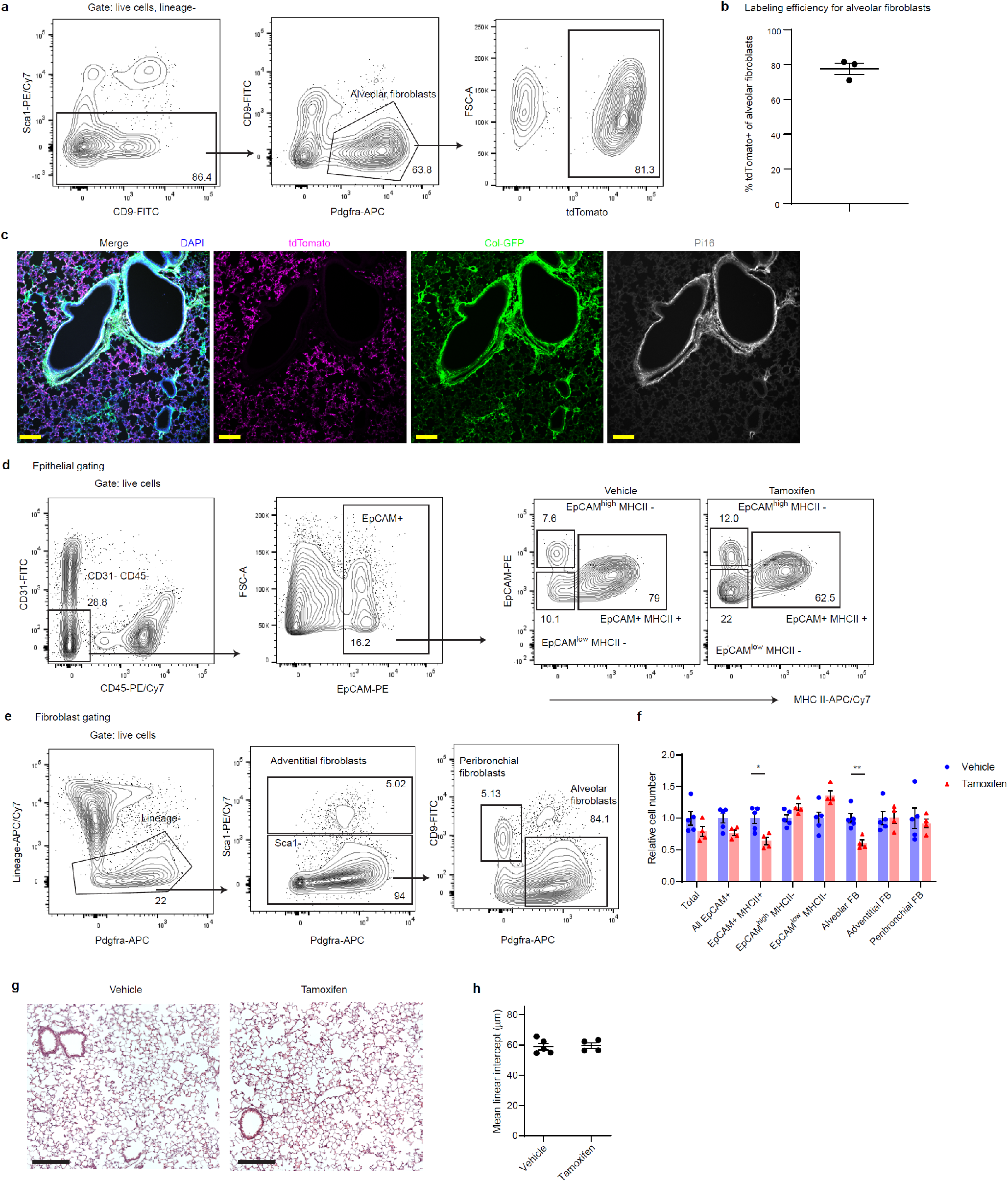
Scube2-CreER specifically labels alveolar fibroblasts, which provide a niche to support AT2 cells. (a) Gating strategy for alveolar fibroblasts. tdTomato+ cells among alveolar fibroblasts are 81.3%. (b) Flow cytometric quantification for percent tdTomato+ of alveolar fibroblasts. n = 3 mice. (c) Pi16 staining of a lung section from Scube2-CreER/Rosa26-tdTomato Col-GFP mouse. tdTomato is shown in magenta. Col-GFP is shown in green. Pi16 is shown in grey. DAPI is shown in blue. (d, e) Gating strategy for alveolar fibroblast ablation experiments with Scub2-CreER/Rosa26-DTA mice. (d) Gating strategy for EpCAM+ subpopulations. (e) Gating strategy for fibroblast subsets. Lineage markers include CD31, CD45, EpCAM, and Ter119. (f) Flow cytometric cell count for each population, normalized to means of vehicle groups. *p = 0.0128, **p = 0.0040 (unpaired two-tailed t-test). (g) Representative images of H&E staining of lung sections from alveolar fibroblast ablation experiments. (h) Quantification of mean linear intercept of alveolar regions. Scale bars, 200 μm (c, g). Data are mean ± SEM. Data are representative of at least two independent experiments.

**Supplementary Fig. 3.**
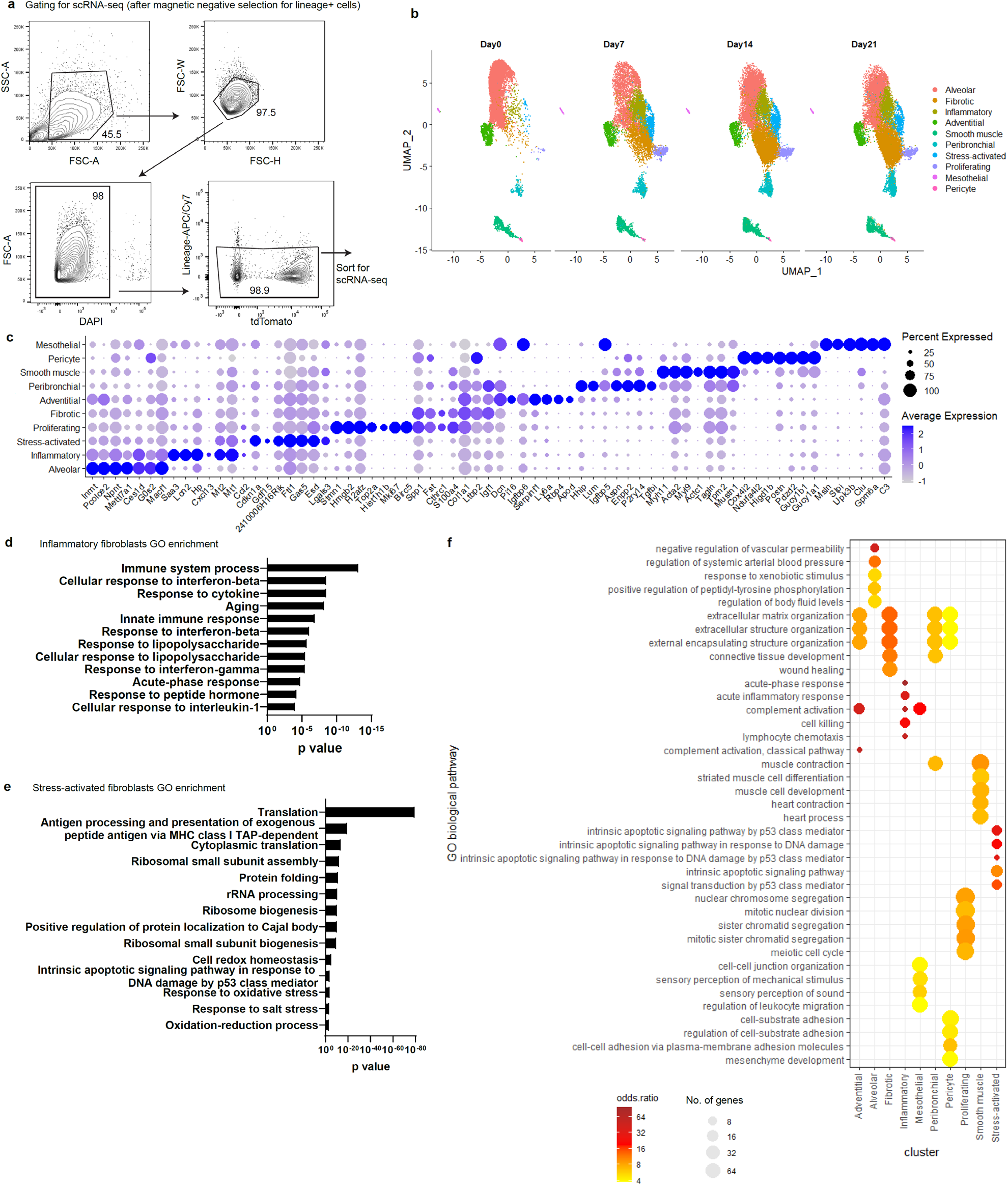
Longitudinal scRNA-seq reveals multiple fibroblast subsets that emerge after lung injury. (a) Gating strategy for purifying lineage (CD31, CD45, EpCAM, Ter119)-mesenchymal cells for scRNA-seq. (b) UMAP plots for cells obtained before (day 0) or at various time points after bleomycin treatment. (c) Dot plots showing top differentially expressed genes for each cluster. (d) GO enrichment analysis for differentially expressed genes of inflammatory fibroblasts. (e) GO enrichment analysis for differentially expressed genes of stress-activated fibroblasts. (f) GO over-representation analysis with Fisher test for all clusters.

**Supplementary Fig. 4.**
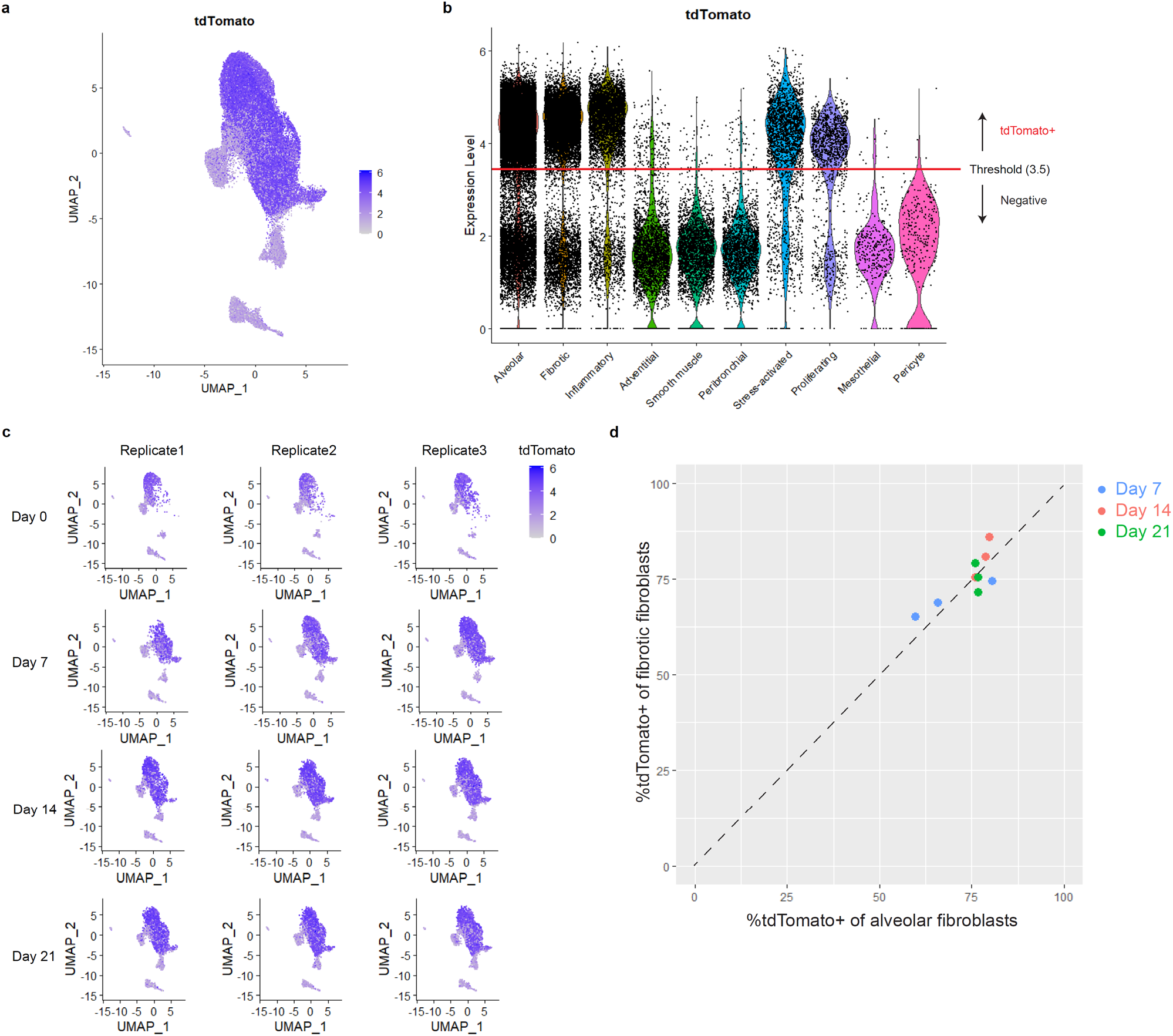
Lineage tracing by scRNA-seq reveals alveolar fibroblasts as origin of multiple pathologic fibroblast subsets. (a) UMAP plot with tdTomato expression. (b) Violin plot for tdTomato shows peaks for tdTomato^low^ or tdTomato^high^ cells. Threshold for tdTomato+ cells was defined as expression level > 3.5. (c) UMAP plots with tdTomato expression split by biological replicates. (d) Plot showing percent tdTomato+ of alveolar fibroblasts (x axis) versus percent tdTomato+ of fibrotic fibroblasts (y axis) for each biological replicate.

**Supplementary Fig. 5.**
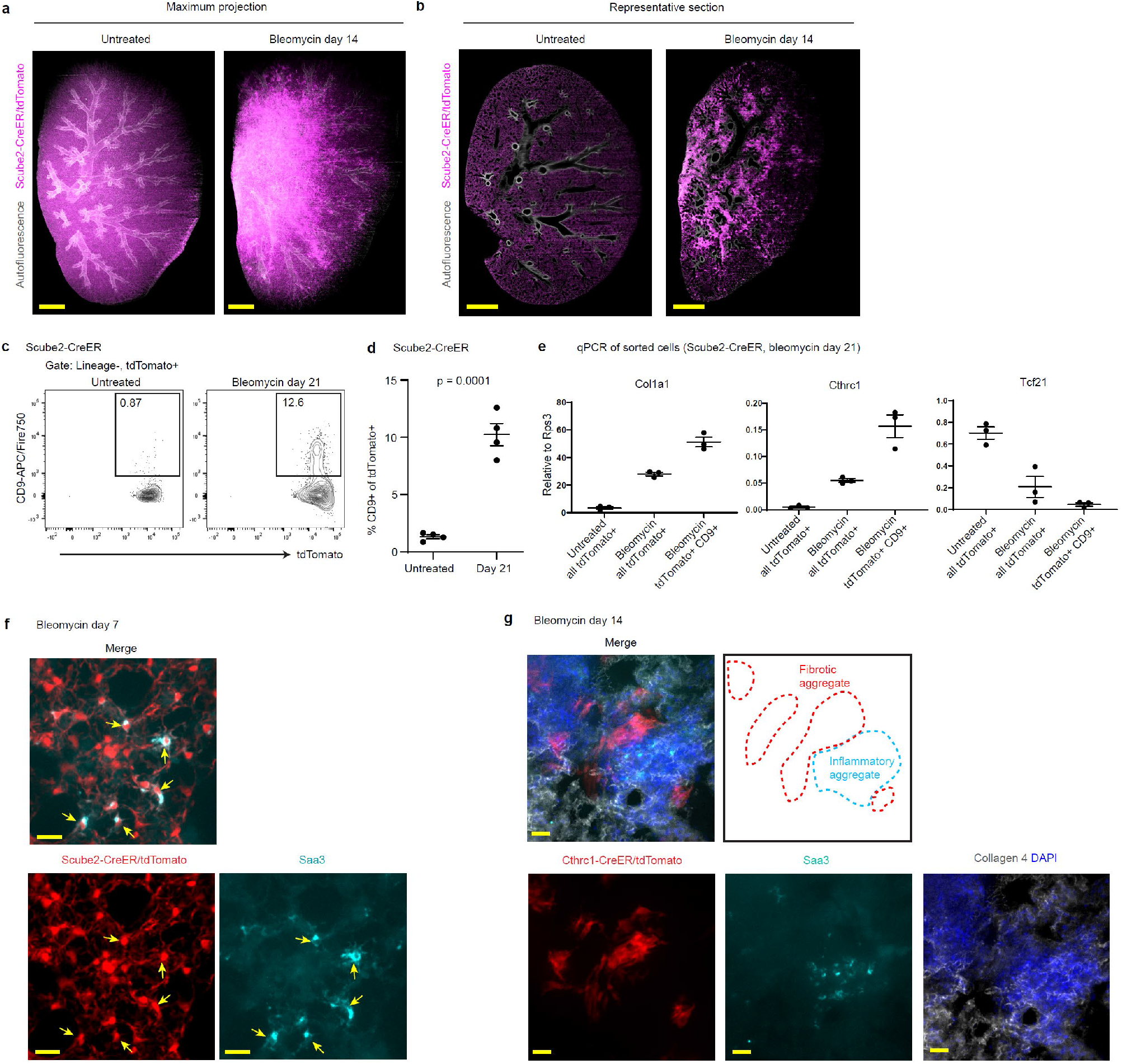
Scube2-CreER-labeled alveolar fibroblasts differentiate into fibrotic or inflammatory fibroblasts after lung injury. (a) Maximum projection of whole lung imaging for untreated or bleomycin day 14 Scube2-CreER/Rosa26-tdTomato mice. (b) Representative optical sections from whole lung imaging. tdTomato is shown in magenta. Autofluorescence is shown in grey (a, b). (c) Flow cytometry plots showing the increase of CD9+ cells among Scube2-CreER-labeled (tdTomato+) cells on day 21 after bleomycin treatment. (d) Flow cytometric quantification of precent CD9+ of tdTomato+ cells. n = 4 mice. Statistical analysis was performed using unpaired two-tailed t-test. (e) qPCR analysis of sorted cells from Scube2-CreER/Rosa26-tdTomato mice. All lineage (CD31, CD45, EpCAM, Ter119)-tdTomato+ cells (untreated or bleomycin day 21) or lineage-tdTomato+ CD9+ cells (bleomycin day 21) were sorted. Y axis is relative expression level to housekeeping gene Rps3. (f) Saa3 staining in sections from Scube2-CreER/Rosa26-tdTomato mice 7 days after bleomycin treatment. Arrows indicate tdTomato and Saa3 double positive cells. (g) Saa3 staining in sections from Cthrc1-CreER/Rosa26-tdTomato mice 14 days after bleomycin treatment with tamoxifen injected on day 8 - 12. tdTomato is shown in red, Saa3 is shown in cyan (f, g). DAPI is shown in blue, Collagen 4 is shown in grey (g). Data are mean ± SEM. Data are representative of at least two independent experiments. Scale bars, 1mm (a, b), 20 μm (f, g).

**Supplementary Fig. 6.**
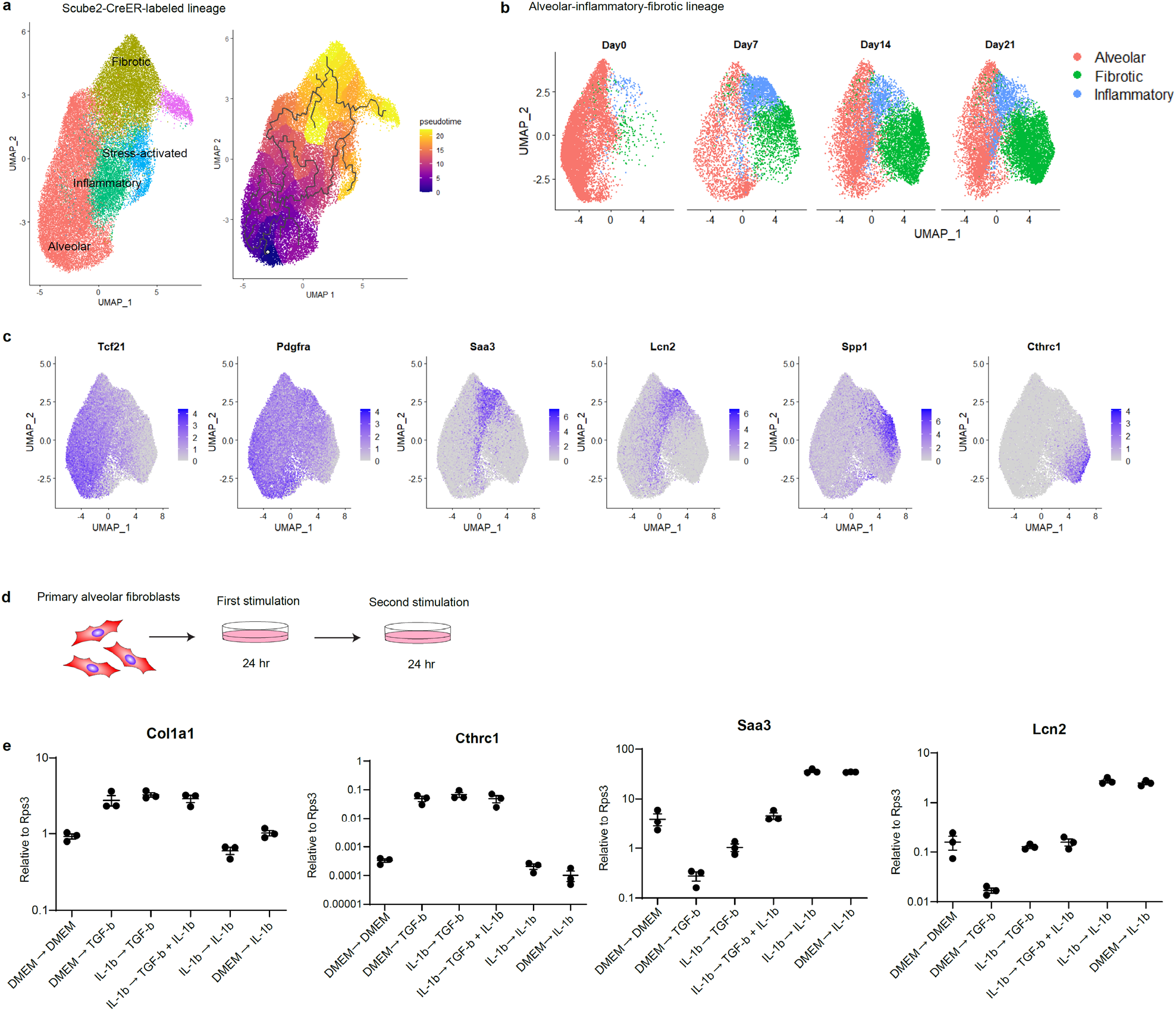
Pseudotime and in vitro analysis suggests that IL-1β and TGF-β regulate sequential differentiation from alveolar into inflammatory and fibrotic fibroblasts. (a) Pseudotime analysis of tdTomato+ clusters suggests that both stress-activated fibroblasts and fibrotic fibroblasts can emerge from inflammatory fibroblasts. (b) UMAP plots of re-clustered alveolar, inflammatory, and fibrotic lineage split by days after bleomycin treatment. (c) Expression of selected markers on UMAP plots. (d) Schematic of sequential cytokine stimulations for primary alveolar fibroblasts. (e) qPCR analysis after sequential cytokine stimulations. Group names indicate (first stimulation) → (second stimulation). DMEM means medium only control. Y axis is relative expression level to housekeeping gene Rps3. Data are representative of two independent experiments

**Supplementary Fig. 7.**
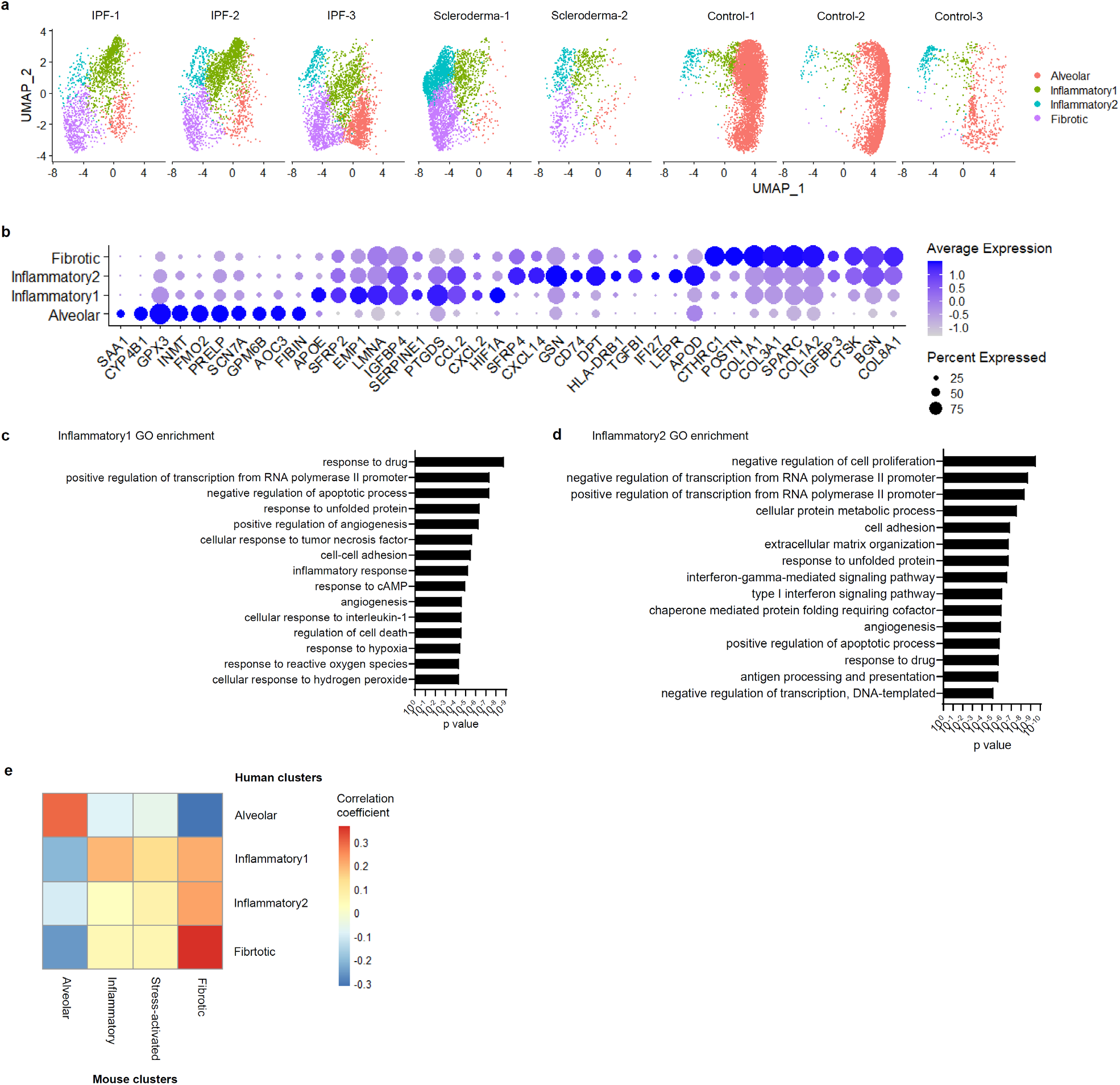
Re-analysis of our publicly available scRNA-seq data from human pulmonary fibrosis reveals inflammatory and fibrotic clusters. (a) UMAP plots of re-clustered pathologic and alveolar clusters shown for individual patients or donors (control). (b) Dot plot showing top differentially expressed genes for each cluster. (c) GO enrichment analysis for differentially expressed genes in inflammatory1 cluster. (d) GO enrichment analysis for differentially expressed genes in inflammatory2 cluster. (e) Heat map showing correlation coefficient of average gene expression from mouse and human emergent clusters.

**Supplementary Fig. 8.**
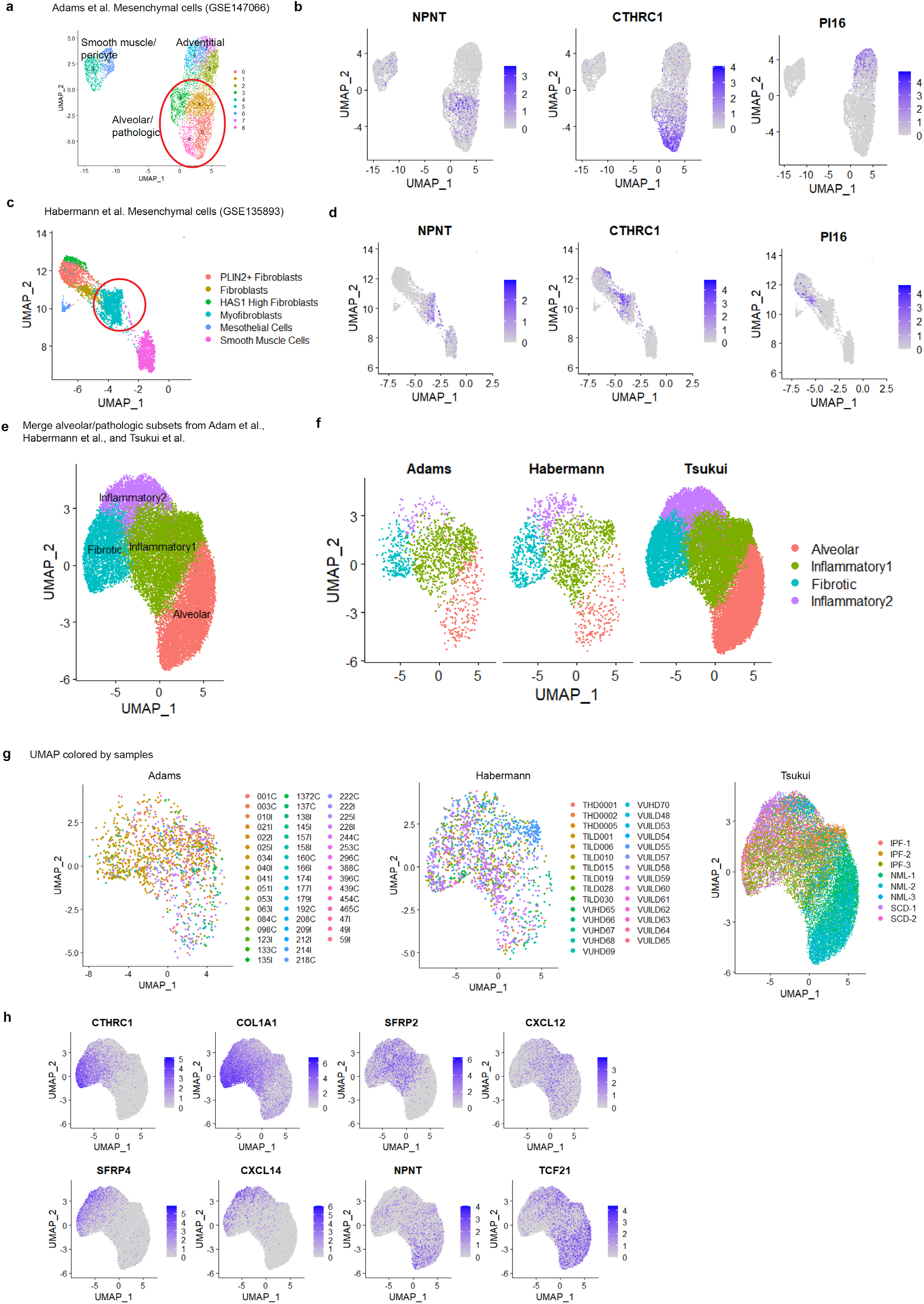
Combined analysis of publicly available scRNA-seq data sets from human pulmonary fibrosis from 3 groups reveals conserved inflammatory and fibrotic clusters. (a) UMAP plot of mesenchymal cells from Adams et al. (b) Expression levels of selected genes on UMAP plot of Adams et al. mesenchymal cells show alveolar and pathologic fibroblast clusters. (c) UMAP plot of mesenchymal cells from Habermann et al. (d) Expression levels of selected genes on UMAP plot of Habermann et al. mesenchymal cells show that “Myofibroblasts” cluster contains alveolar and pathologic fibroblasts. (e) UMAP plot after combining alveolar and pathologic fibroblasts from Adams et al., Habermann et al., and Tsukui et al. (f) UMAP plots of combined data split by original data set. (g) UMAP plots shown for each data set and colored by samples. (h) Expression levels of selected markers for fibrotic (COL1A1, CTHRC1), inflammatory1 (SFRP2, CSCL12), inflammatory2 (SFRP4, CXCL14) and alveolar fibroblasts (NPNT, TCF21) on UMAP plots.

## Reference

1. Plikus, M. V. et al. Fibroblasts: Origins, definitions, and functions in health and disease. Cell 184, 3852–3872 (2021).

2. Hinz, B. & Lagares, D. Evasion of apoptosis by myofibroblasts: a hallmark of fibrotic diseases. Nat Rev Rheumatol 16, 11–31 (2020).

3. Henderson, N. C., Rieder, F. & Wynn, T. A. Fibrosis: from mechanisms to medicines. Nature 587, 555–566 (2020).

4. Tsukui, T. et al. Collagen-producing lung cell atlas identifies multiple subsets with distinct localization and relevance to fibrosis. Nat Commun 11, 1920 (2020).

5. Buechler, M. B. et al. Cross-tissue organization of the fibroblast lineage. Nature 593, 575–579 (2021).

6. Ruiz-Villalba, A. et al. Single-Cell RNA Sequencing Analysis Reveals a Crucial Role for CTHRC1 (Collagen Triple Helix Repeat Containing 1) Cardiac Fibroblasts After Myocardial Infarction. Circulation 142, 1831–1847 (2020).

7. Melms, J. C. et al. A molecular single-cell lung atlas of lethal COVID-19. Nature 595, 114–119 (2021).

8. Pakshir, P. et al. The myofibroblast at a glance. Journal of Cell Science 133, jcs227900 (2020).

9. Friedman, S. L., Sheppard, D., Duffield, J. S. & Violette, S. Therapy for Fibrotic Diseases: Nearing the Starting Line. Science Translational Medicine 5, 167sr1–167sr1 (2013).

10. Narvaez del Pilar, O., Gacha Garay, M. J. & Chen, J. Three-axis classification of mouse lung mesenchymal cells reveals two populations of myofibroblasts. Development 149, dev200081 (2022).

11. Barratt, S. L., Creamer, A., Hayton, C. & Chaudhuri, N. Idiopathic Pulmonary Fibrosis (IPF): An Overview. J Clin Med 7, 201 (2018).

12. McGowan, S. E. & Torday, J. S. The pulmonary lipofibroblast (lipid interstitial cell) and its contributions to alveolar development. Annu Rev Physiol 59, 43–62 (1997).

13. Barkauskas, C. E. et al. Type 2 alveolar cells are stem cells in adult lung. J Clin Invest 123, 3025–3036 (2013).

14. Hasegawa, K. et al. Fraction of MHCII and EpCAM expression characterizes distal lung epithelial cells for alveolar type 2 cell isolation. Respiratory Research 18, 150 (2017).

15. Croft, A. P. et al. Distinct fibroblast subsets drive inflammation and damage in arthritis. Nature 570, 246–251 (2019).

16. Öhlund, D. et al. Distinct populations of inflammatory fibroblasts and myofibroblasts in pancreatic cancer. Journal of Experimental Medicine 214, 579–596 (2017).

17. Boyd, D. F. et al. Exuberant fibroblast activity compromises lung function via ADAMTS4. Nature 587, 466–471 (2020).

18. Korsunsky, I. et al. Cross-tissue, single-cell stromal atlas identifies shared pathological fibroblast phenotypes in four chronic inflammatory diseases. Med 0, (2022).

19. Cao, J. et al. The single-cell transcriptional landscape of mammalian organogenesis. Nature 566, 496–502 (2019).

20. Elyada, E. et al. Cross-Species Single-Cell Analysis of Pancreatic Ductal Adenocarcinoma Reveals Antigen-Presenting Cancer-Associated Fibroblasts. Cancer Discovery 9, 1102–1123 (2019).

21. Adams, T. S. et al. Single-cell RNA-seq reveals ectopic and aberrant lung-resident cell populations in idiopathic pulmonary fibrosis. Science Advances 6, eaba1983 (2020).

22. Habermann, A. C. et al. Single-cell RNA sequencing reveals profibrotic roles of distinct epithelial and mesenchymal lineages in pulmonary fibrosis. Science Advances 6, eaba1972 (2020).

23. Valenzi, E. et al. Single-cell analysis reveals fibroblast heterogeneity and myofibroblasts in systemic sclerosis-associated interstitial lung disease. Annals of the Rheumatic Diseases 78, 1379–1387 (2019).

24. Dominguez, C. X. et al. Single-Cell RNA Sequencing Reveals Stromal Evolution into LRRC15+ Myofibroblasts as a Determinant of Patient Response to Cancer Immunotherapy. Cancer Discovery 10, 232–253 (2020).

25. Binks, A. P., Beyer, M., Miller, R. & LeClair, R. J. Cthrc1 lowers pulmonary collagen associated with bleomycin-induced fibrosis and protects lung function. Physiol Rep 5, e13115 (2017).

26. Li, J. et al. Autocrine CTHRC1 activates hepatic stellate cells and promotes liver fibrosis by activating TGF-β signaling. EBioMedicine 40, 43–55 (2019).

27. Yata, Y. et al. DNase I-hypersensitive sites enhance alpha1(I) collagen gene expression in hepatic stellate cells. Hepatology 37, 267–276 (2003).

28. Henderson, N. C. et al. Targeting of αv integrin identifies a core molecular pathway that regulates fibrosis in several organs. Nat Med 19, 1617–1624 (2013).

29. Madisen, L. et al. A robust and high-throughput Cre reporting and characterization system for the whole mouse brain. Nat Neurosci 13, 133–140 (2010).

30. Hao, Y. et al. Integrated analysis of multimodal single-cell data. Cell 184, 3573–3587.e29 (2021).

31. McGinnis, C. S., Murrow, L. M. & Gartner, Z. J. DoubletFinder: Doublet Detection in Single-Cell RNA Sequencing Data Using Artificial Nearest Neighbors. cels 8, 329–337.e4 (2019).

32. Haghverdi, L., Lun, A. T. L., Morgan, M. D. & Marioni, J. C. Batch effects in single-cell RNA-sequencing data are corrected by matching mutual nearest neighbors. Nat Biotechnol 36, 421–427 (2018).

33. Street, K. et al. Slingshot: cell lineage and pseudotime inference for single-cell transcriptomics. BMC Genomics 19, 477 (2018).

34. Susaki, E. A. et al. Versatile whole-organ/body staining and imaging based on electrolyte-gel properties of biological tissues. Nat Commun 11, 1982 (2020).

35. Crowley, G. et al. Quantitative lung morphology: semi-automated measurement of mean linear intercept. BMC Pulm Med 19, 206 (2019).

